# Rewiring of RNA methylation by the oncometabolite fumarate in renal cell carcinoma

**DOI:** 10.1101/2023.04.10.536262

**Authors:** Christina M. Fitzsimmons, Mariana D. Mandler, Judith C. Lunger, Dalen Chan, Siddhardha S. Maligireddy, Alexandra C. Schmiechen, Supuni Thalalla Gamage, Courtney Link, Lisa M. Jenkins, Daniel R. Crooks, Jordan L. Meier, W. Marston Linehan, Pedro J. Batista

## Abstract

Metabolic reprogramming is a hallmark of cancer that facilitates changes in many adaptive biological processes. Mutations in the tricarboxylic acid (TCA) cycle enzyme fumarate hydratase (FH) lead to fumarate accumulation and cause hereditary leiomyomatosis and renal cell cancer (HLRCC). HLRCC is a rare, inherited disease characterized by the development of non-cancerous smooth muscle tumors of the uterus and skin, and an increased risk of a highly metastatic and aggressive form of kidney cancer. Fumarate has been shown to inhibit 2-oxyglutarate-dependent dioxygenases (2OGDDs) involved in the hydroxylation of HIF1α, as well as in DNA and histone demethylation. However, the link between fumarate accumulation and changes in RNA post-transcriptional modifications has not been defined. Here, we determine the consequences of fumarate accumulation on the activity of different members of the 2OGDD family targeting RNA modifications. By evaluating multiple RNA modifications in patient-derived HLRCC cell lines, we show that mutation of FH selectively alters the activity of demethylases acting upon N6-methyladenosine (m^6^A), while the demethylase acting upon N1-methyladenosine (m^1^A) and 5-formylcytosine (f^5^C) in mitochondrial RNA are unaffected. The observation that metabolites modulate specific subsets of RNA-modifying enzymes offers new insights into the intersection between metabolism and the epitranscriptome.

## INTRODUCTION

Cellular plasticity and identity depend upon metabolic and environmental cues, and metabolic rewiring is essential for commitment to specific cellular lineages during all stages of development (1). The tricarboxylic acid (TCA) cycle is a central metabolic pathway used by organisms to generate energy, in the form of ATP, as well as building blocks for protein, nucleic acid, and fatty acid synthesis (2). Cellular metabolic reprogramming is an important mechanism by which cells rewire their metabolism to promote proliferation. This process is frequently observed in tumors, and is considered a hallmark of cancer (3, 4). Aerobic glycolysis, commonly referred to as the Warburg effect, is characterized by decreased ATP synthesis and increased aerobic metabolism of glucose to lactate. Mutations in TCA cycle enzymes that disrupt oxidative phosphorylation are a known driver of metabolic rewiring and have been linked to specific cancer subtypes, such as gliomas, acute myeloid leukemia, gastrointestinal stromal tumors, and paragangliomas (5–10). To date, mutations associated with oncogenesis have been described for four TCA enzymes: isocitrate dehydrogenase 1 and 2 (IDH1 and IDH2) (11–13), succinate dehydrogenase (SDH) and fumarate hydratase (FH). Gain-of-function mutations in IDH1 and IDH2 lead to the *de novo* production of 2-hydroxyglutarate (2HG) (5, 6). Conversely, loss of SDH or FH activity results in the accumulation of succinate and fumarate, respectively (7, 8, 14–16). Critically, aberrant accumulation of these metabolites can disrupt the activity of cellular enzymes, leading to changes in gene expression and signaling, thereby promoting tumorigenesis. Loss of function mutations in FH have been identified in individuals with the familial cancer syndrome Hereditary Leiomyomatosis and Renal Cell Carcinoma (HLRCC) (15–17). These patients are at risk for the development of cutaneous and uterine leiomyomas as well as kidney tumors. These renal tumors readily metastasize, leading to a highly aggressive form of renal cell carcinoma (18).

One group of enzymes that is sensitive to changes in TCA cycle metabolites, including fumarate accumulation, is the 2-oxoglutarate-dependent dioxygenase (2OGDD; also known as alpha-ketoglutarate (αKG)-dependent dioxygenases) family (11, 12). Enzymes in this family catalyze hydroxylation reactions on lipids, proteins, and nucleic acids (19). Critically, the activity of these enzymes is dependent upon the availability of iron, oxygen, and the ratio of αKG to related TCA cycle metabolites such as succinate and fumarate (20, 21). Aberrant accumulation of 2HG, succinate, or fumarate can disrupt the activity of 2OGDDs, leading to changes in gene expression programs and promoting tumorigenesis. Previous work has demonstrated that high levels of fumarate can upregulate the transcription factor Hypoxia Inducible Factor (HIF) by inhibiting HIF prolyl hydroxylase activity (22). Additionally, accumulation of fumarate has been shown to promote hypermethylation of DNA and histones (23, 24) and drive an epithelial-to-mesenchymal (EMT) phenotypic switch in HLRCC (25).

While previous studies have focused on the effect of metabolic changes on DNA and chromatin demethylation, less is known about the impact of fumarate accumulation on 2OGDDs that act on RNA substrates. The AlkB proteins are a highly conserved family of 2OGDDs with nine human homologs (ALKBH1-8 and FTO). Five of these members have been demonstrated to act on RNA (ALKBH1, ALKBH3, ALKBH5, ALKBH7, and FTO) (reviewed in (26, 27)). These enzymes catalyze the removal of RNA modifications and can act on a diverse set of modified nucleobases and types of RNA including messenger RNA (mRNA), transfer RNA (tRNA), and ribosomal RNA (rRNA). Recent studies have revealed roles for several ALKBH family members in cancer, where they can function either as oncogenes or tumor suppressors in a cell-type specific fashion (28–30). Considering the potential for ALKBH enzymes to serve as a link between metabolism and gene expression, we set out to define the consequences of fumarate accumulation in HLRCC on the levels of RNA modifications that are substrates of 2OGDD enzymes. Here we report that mutation of FH alters the activity of demethylases acting up N6-methyladenosine (m^6^A), while demethylases acting upon N1-methyladenosine (m^1^A) and 5-formylcytosine (f^5^C) are unaffected. The observation that certain 2OGDD enzymes are more sensitive than others to fumarate accumulation offers new insights into the intersection of metabolism and gene regulation.

## MATERIALS & METHODS

### General cell culture

The parental UOK262 (15) and HK2 (ATCC CRL-2190) were grown in heat-inactivated Dulbecco’s modification of Eagle’s medium (DMEM; Gibco 10313021) supplemented with 10% heat-inactivated fetal bovine serum (FBS; Sigma Aldrich #F4135), 2% glutamax, (ThermoFisher #35050061) and 1% penicillin-streptomycin (PenStrep; ThermoFisher #15140122). HEK-293T cells (ATCC CRL-11268) were grown in DMEM supplemented with 10% fetal bovine serum, 1% glutamax, and 1% PenStrep. Cells were grown at 37°C and 5% carbon dioxide and were split every 2-3 days at approximately 80% confluency. All cell lines were routinely tested for mycoplasma contamination by the MycoAlert Mycoplasma Detection Kit (Lonza #LT07-318).

### Generation of isogenic UOK cell lines

The stable UOK262 isogenic cell lines were generated by lentivirus. Briefly, the FH gene was cloned into the pLEX_305 lentiviral backbone where we swapped the puromycin resistance for geneticin resistance. The pLEX_305 vector was a gift from David Root (Addgene plasmid #41390). Viral particles were generated using 2nd generation lentiviral system plasmids (31). Briefly, 4.5 µg of Δ8.9 plasmid, 1.5 µg of vsv-g plasmid (gifts from Howard Chang), 6 µg of pLEX_305 vector plasmid, 600 µL of Opti-MEM (ThermoFisher #31985062) and 36 µL of Fugene (Promega #E2691) were mixed and incubated at room temperature for 15 min before being added to a 10-cm plate containing 5 million HEK-293T cells. The following day, the media was changed on the 10-cm plate. After 2 days, media was removed and concentrated using lenti-X concentrator solution (Takara Bio #631232). Concentrated viral particles were distributed into single-use aliquots and stored at −80°C until use. Viral aliquots were titrated using the Lenti-X qRT-PCR Titration Kit (Clonetech #631235). For cell line transduction of the UOK cells, a range of viral dilutions was added to the cells. The following day, media was changed, and cells were selected with geneticin (G418; ThermoFisher #10131035) for 4-5 days. A mock viral-treated plate containing UOK262 cells was used as a control for G418 selection. Introduction of FH-FLAG was confirmed by qPCR and Western blot, and WT and Mutant cell lines were paired based on FH expression.

### qPCR

Total RNA was collected from 1-2 million UOK262 cells using Trizol reagent (ThermoFisher #15596026) according to the manufacturer’s protocol. Following separation of the aqueous and organic phases, the aqueous phase was removed and purified with a Qiagen RNeasy Mini kit (#74004) according to the manufacturer’s protocol. RNA was eluted in 100 µL of nuclease-free water and RNA concentration and purity were determined by nano-volume spectrophotometric analysis (DeNovix). Reverse Transcription (RT) was performed with 0.5 to 1.0 µg of total RNA input material in 20 µL of total volume using SuperScript IV Vilo reverse transcriptase (ThermoFisher #11754050) and following the manufacturer’s protocol for thermal cycling: 25°C for 10 min; 50°C for 10 min; 85°C for 5 min; 4°C hold. A control reaction containing the no reverse-transcriptase solution was also prepared. Upon completion of the reaction, cDNA was diluted to a final volume of 100 µL in nuclease-free water. In some experiments, the reverse-transcribed cDNA was stored overnight at 4°C. The RT reaction was primed with random nucleotide hexamers present in the Vilo master mix.

Quantitative real-time polymerase chain reaction (qPCR) reactions were setup manually in 96-well plates using either Sybr Green chemistry (25 µL reactions using Applied Biosystems Power SYBR green PCR mix #4367659) or hydrolysis probe chemistry (20 µL reactions Taqman probes, ThermoFisher) and the following the manufacturers’ protocols. Sybr Green chemistry reactions: 12.5 µL 2X Sybr Green Master Mix, 3 µL of cDNA, 2 µL of 10 mM primer mix, 7.5 µL nuclease-free water. Primers for the Sybr Green qPCR reactions (Table S1) were ordered from IDT with no modifications and standard desalting. Hydrolysis probe chemistry reactions: 1 µL 20X Taqman gene expression probe; 10 µL 2X Taqman Gene Expression Fast Advanced Master Mix (ThermoFisher #4444557), 4 µL cDNA template, 5 µL Nuclease-free water.

Amplification was measured on a QuantStudio3 RT-qPCR thermal cycler using the recommended parameters for each chemistry. Sybr Green reactions were performed in standard run mode with the following parameters: 95°C for 10 min, 40 cycles of 95°C for 15 sec, 60°C for 60 sec. Melting curves were measured from 65°C to 95°C in steps of 0.5°C. Hydrolysis probe reactions were performed in fast run mode with the following parameters: 50°C for 2 min, 95°C for 20 sec, 40 cycles of 95°C for 1 sec, 60°C for 20 sec. Hydrolysis probes (Table S1) were ordered from ThermoFisher Scientific. To test for contamination of the RNA, we performed control reactions to omit the SuperScript IV. To test for contamination of the PCR reagents, we performed “no-template controls” (NTC). mRNA values were normalized relative to reference genes GAPDH, Actin, or PP1A. The standard comparative cycle threshold method was used for quantification. All reverse transcription reactions (biological replicates) and qPCR reactions (technical replicates) were performed in a minimum of triplicate. Data are plotted as mean ± sd in either GraphPad Prism (v 9.0) or R-studio (v 3.6.3).

### Western Blot

Cell pellets from 100, 000-300, 000 UOK262 cells were lysed in RIPA buffer (50 mM Tris, pH 7.4, 150 mM NaCl, 1% NP-40, 0.1% SDS, 0.5% Sodium deoxycholate, supplemented with a protease inhibitor cocktail tablet (Millipore Sigma # 11697498001). Lysates were harvested by centrifugation at 21, 000 x *g* for 30 min at 4°C. Protein was quantified by bicinchoninic acid (BCA) assay (ThermoFisher #23227). Protein (between 10 and 20 ug) was resolved by sodium dodecyl sulfate–polyacrylamide gel electrophoresis (SDS-PAGE; ThermoFisher #NP0335) in 1X MOPS buffer (40 mM 3-(N-morpholino) propanesulfonic acid (MOPS), 10 mM sodium acetate, 1 mM EDTA), transferred to nitrocellulose membranes (Li-COR #926-31092), and membranes were blocked in Intercept (TBS) Blocking Buffer (LI-COR #927-60001). Membranes were incubated overnight with primary antibodies at 4°C with gentle rocking. The following morning, membranes were rinsed 3X with phosphate buffered saline supplemented with Tween-20 (PBS-T; 1.37 M NaCl, 27 mM KCl, 100 mM Na_2_HPO_4_, 18 mM KH_2_PO_4_, 0.5% Tween-20) and incubated with fluorophore-conjugated species-specific secondary antibodies (LI-COR) for 1 hour at 20°C with gentle rocking. Fluorescence was detected on a LI-COR Odyssey instrument and analyzed using the LI-COR ImageStudio software. A list of all primary and secondary antibodies as well as the dilution factors used is provided in Table S2.

### Subcellular Fractionation

For subcellular fractionation, the procedure was adapted from previously described protocols (32). Briefly, 1.5 million trypsinized cells were washed 2X with 5 mL of PBS and re-suspended thoroughly in 600 μL HEPES-Sucrose-Ficoll-Digitonin solution (HSFD, 20 mM HEPES-KOH, 6.25% Ficoll, 0.27 M sucrose, 3 mM CaCl_2_, 2 mM MgCl_2_. pH7.4) with freshly added digitonin and proteinase inhibitors and lysed on ice for 10 min. Following lysis, cells were centrifuged at 1, 000 x *g* for 3 min and the supernatant was removed and centrifuged again at 15, 000 x *g* for 10 min to generate the cytosol fraction. The pellet of nuclei was washed with HSFD buffer while the supernatant was collected as the washed fraction. The pellet was lysed by sonication in 100 µL of RIPA buffer. Following sonication, fraction was incubated on ice for 30 min, and then centrifuged to 15, 000 x *g* for 10 min. After centrifugation, the supernatant was collected as the nuclear extract fraction. The concentration of protein from each fraction was quantified by BCA Assay (ThermoFisher #23227). Protein (between 10 and 20 µg) was resolved by SDS-PAGE and proteins of interest were detected as above described.

### FH and MDH Native In-gel Activity Assay

Cell collected from 1 T75 flask (approx. 1-2 million) were pelleted and frozen until use. Frozen cell pellets were lysed by sonication on ice in 200 µL pre-chilled Lysis Buffer (10 mM Tris, pH 8.0, 0.1% Triton X-100, 3 mM KCl, 3 mM MgCl_2_, 3 mM sodium citrate) supplemented with an EDTA-free protease inhibitor tablet (SigmaAldrich #04693116001). Lysates were clarified by centrifugation at 4°C for 30 min at 21, 000 x *g*. Supernatant was removed and protein concentration was quantified by BCA assay (ThermoFisher #23227) according to the manufacturer’s protocol. 20 µg of protein sample was loaded onto 7% Tris-Acetate gels (ThermoFisher #EA03552) with ice-cold 1X Tris-glycine buffer and run at 125V and 4°C for approximately 1 hr. Upon completion, gels were washed gently with deionized water for 5 minutes at 20°C with gentle rocking. After 5 min, 10 mL of fumarate assay buffer (0.1 M Tris, pH 7.4, 1 mg/mL 3-(4, 5-dimethylthiazol-2-yl)-2, 5-diphenyltetrazolium bromide (MTT; Sigma Aldrich #M2128), 1 mM NAD+, 20 µL malic dehydrogenase (SigmaAldrich #M2634), 0.182 mg/mL phenazine methosulfate (PMS; Sigma Aldrich #P9625) and 5 mM sodium fumarate (SigmaAldrich #F1506)) or malic dehydrogenase assay buffer (0.1M Tris, pH 7.4, 1 mg/mL MTT, 0.182 mg/mL PMS, 5 mM L-malate (Sigma Aldrich #112577) were added to the tray containing the gels. Gels were incubated for 5-15 min in a heated rocker oven at 37°C until colorimetric bands appeared in the native gel. Gels were destained in ddH_2_O for 5-15 min and scanned on an Epson Perfection V700 Scanner in transmitted color mode.

### Immunohistochemistry

Five-micron thick formalin-fixed paraffin-embedded sections were cut from UOK262 tumor xenografts and a human renal cortex specimen. Immunohistochemistry staining was performed by VitroVivo Biotech (Rockville, MD) using standardized methodologies, using either anti-ALKBH5 (1:250) or anti-FTO (1:800). Images were captured using an AxioScan.Z1 Slide Scanner (Zeiss, Oberkochen, DE). A list of antibodies is provided in Table S2.

### Immunofluorescence

Adherent cells were grown on ethanol-sterilized glass coverslips previously coated with Poly-D-Lysine Solution (ThermoFisher #A3890401). Cells were fixed with 4% paraformaldehyde (PFA; ThermoFisher #28908) for 10 min and then permeabilized with 0.3% Triton X-100 for 5 min. Blocking was carried out for 1 hr with PBS containing 5% FBS. Primary antibodies were incubated overnight at 4°C in blocking solution. Coverslips were washed gently for 15 minutes with PBS and then incubated for 1 hr at 20°C with secondary antibodies. Coverslips were gently washed using STAR (v 2.7.6a) (35, 36). Duplicates were marked and removed with Picard’s MarkDuplicates (v2.25.0). Sample counts were extracted using htseq-count function (v 0.9.1) (37) and differential gene expression was performed with the DeSeq2 package (38) in R (v 3.6.3)

### m^6^A-immunoprecipitation

Total RNA was collected from 1-2 million UOK262 cells using Trizol reagent (ThermoFisher #15596026) according to the manufacturer’s protocol. Following separation of the aqueous and organic phases, the aqueous phase was removed and purified with a Qiagen RNA Mini kit (#74004) according to the manufacturer’s protocol. RNA was eluted in 100 µL of nuclease-free water and RNA concentration and purity were determined by nano-volume spectrophotometric analysis (DeNovix). mRNA was polyA selected through two rounds of selection with oligo d(T)_25_ magnetic beads 3X with PBS. After washing, DNA was counterstained with DAPI (ThermoFisher #62248) and coverslips were mounted with ProLong™ Glass Antifade Mountant (ThermoFisher #P36980). Images were acquired using a Nikon SoRa spinning disk microscope equipped with Photometrics BSI sCMOS camera and 60x apochromat TIRF (N.A. 1.49) oil immersion lens and were analyzed using FIJI (33). A list of antibodies is provided in Table S2.

### RNA sequencing

Total RNA was extracted from cells using Trizol Reagent (ThermoFisher #15596018) according to the manufacture’s protocol. Following phase separation, the aqueous phase was applied to a RNeasy Mini Kit column (Qiagen #74106) and RNA was purified according to kit instructions. RNA-seq libraries were prepared using Illumina TruSeq Stranded Total RNA Library Prep Kit v2 (Illumina #RS-122-2001 and #RS-122-2002) and sequenced on NextSeq 500 using paired-end sequencing (2 x 75). Read quality was checked with FastQC (v 0.11.9). Reads of the samples were trimmed for adapters using cutadapt (v 2.5) (34) and aligned against a costume genome of repetitive sequences before alignment with the hg38 reference genome (gencode v35) using Spliced Transcripts Alignment to a Reference (STAR v2.7.6a (35, 36)). PCR duplicates were marked and removed with Picard’s MarkDuplicates (v 2.25.0). Sample counts were obtained with htseq-count (v 0.9.1) (37) and differential gene expression was analyzed with DeSeq2 package (38) in R (v 3.6.3)

### Analysis of Previously Published HLRCC RNA-seq data

HLRCC patient RNA-seq data generated by a previous study (39) was used to examine changes in gene expression between metastatic and primary tumor sites. Paired fastq data was downloaded from SRA (GSE157256) using sratoolkit and verified based on md5 hash. Adapters were trimmed from the fastq files using cutadapt (v 2.5) (34) before alignment with the hg38 reference genome (gencode v35) (NEB #S1419S). Following polyA selection, 5 µg of mRNA was fragmented using ZnCl_2_ fragmentation reagent (Life Technologies #AM8740) to a size of approximately 100 nucleotides. Fragmented mRNA was purified using Zymo Research RNA Clean and Concentrator Kit (Zymo Research #R1013). For m^6^A-immunoprecipitation (m^6^A-IP) both proteinA (ThermoFisher #10008D) and proteinG beads (ThermoFisher #1009D) were bound overnight at 4°C with tumbling end-over-end to rabbit polycolonal anti-m^6^A (Synaptic Systems #202-003). m^6^A-IP was performed in 200 µL of IP buffer (150 mM NaCl, 0.1% NP-40, 10 mM Tris, pH 7.5) with 1 µL of RNAseOUT (ThermoFisher #10777019) for 3 hours tumbling end-over-end at 4°C. mRNA was first bound to proteinG beads and 5% of fragmented material was reserved for “input” samples during library construction. Bead-bound antibody-RNA complexes were recovered on a magnetic stand and washed 2X with IP buffer, 2X with high-salt buffer (10 mM Tris, pH 7.5, 500 mM NaCl, 0.001% NP-40) 2X with low-salt buffer (10 mM Tris, pH 7.5, 50 mM NaCl, 0.001% NP-40) and 1X with IP buffer. After all washes, RNA was eluted from the proteinG beads with buffer RLT (Qiagen #74004) and cleaned on a RNeasy spin column and eluted in water. Eluted RNA was added to proteinA-antibody conjugated beads in IP buffer, and the binding, wash, and elution steps were repeated. After washing proteinA beads, the final elution was performed in 100 µL of nuclease-free water. Samples were dried by vacuum freeze drying to a volume < 10 µL.

RNA 3′-ends were dephosphorylated in a 10 µL reaction with 2 µL T4 PNK (NEB; #M0201S) 1 µL of FastAP Thermosensitive Alkaline Phosphatase (ThermoFisher #EF0651), and 1 µL of RiboLock (ThermoFisher #EO0381) for 60 min in dephosphorylation buffer (70 mM Tris pH 6.5, 10 mM MgCl_2_, 1 mM DTT) with shaking. After 60 minutes, the components of the 3′-adaptor reaction were added (1 µL pre-adenylated and 3′-adaptor, 1 µL of 10X RNA ligase Buffer, 1 µL of T4 RNA Ligase (NEB #M0437M), 1 µL of 100 mM DTT, and 6 µL of PEG 8000) and the reaction was incubated overnight with shaking at 16°C.

The following day, excess linker was removed by adding of 2 µL of 10X deadenylation buffer, 1 µL 5′deadenylase (NEB #M0331S), 1 µL of RiboLock (ThermoFisher #EO0381) and 1 µL of RecJF (NEB #M0264S). Reactions were incubated for 1 hr at 37°C. Samples were cleaned up with Zymo Research RNA Clean and Concentrator Kit (Zymo Research #R1013). RNA was reverse transcribed with Superscript IV reverse transcriptase (ThermoFisher #18090050) according to the manufacturer’s protocol. cDNA was purified by Streptavidin C1 magnetic beads (ThermoFisher #65002) in Biotin Buffer (100 mM Tris, pH 7.0, 1 M NaCl, 0.1% Tween-20, 1 mM EDTA). The reaction was allowed to bind for 30 min at 20°C with end-over-end tumbling. After 30 min, the reaction was applied to a magnetic stand and reactions were washed 2X with High Stringency Buffer (15 mM Tris-HCl, pH 7.5, 5 mM EDT, 1% Triton X-100, 1% sodium deoxycholate, 0.001% SDS, 120 mM NaCl, 25 mM KCl) and 2X with PBS. Samples were resuspended in 2 x 6 µL nuclease-free water and were boiled from the beads for 5 min at 95°C. cDNA was circularized in at 15 µL reaction containing 12 µL cDNA, 0.5 µL circLigase II (Lucigen # CL9025K) 1.5 µL 10X circLigase buffer, 0.75 µL of 50 mM MnCl_2_ and nuclease-free water to 15 µL. Reactions were incubated for 2 hr at 60°C and purified by Ampure XP beads (Beckman #A63881). Libraries were PCR amplified in a 30 µL reaction containing 14 µL circularized library, 15µL Phusion HF 2X master mix (NEB #M0531L), 0.75 µL of 20 µM P3/P6 tall primer mix and 0.25 µL of 25X Sybr Green (ThermoFisher #S7563). Libraries were amplified for 8-12 cycles as determined by RT-qPCR on a QuantStudio3 RT-qPCR thermal cycler. Amplified libraries were cleaned up using Ampure XP beads and eluted in 10 µL of nuclease-free water. Three additional cycles of PCR were performed in 20 µL total volume containing 10 uL Phusion HF 2X master mix and 250 nM of P3/P6 Solexa primer mix. All primers are listed in Table S1.

Library cleanup was performed using Zymo DNA Cleanup kit (Zymo #D4014) and the manufacturer’s protocol to remove dNTPs and to concentrate the volume. Libraries were then size selected by gel purifying DNA on a 10% TBE gel and cleanup was performed using Zymo DNA Cleanup kit (Zymo # D4014) and the manufacturer’s protocol. Library QC was performed on a DNA High Sensitivity D1000 Tape Station (Agilent) and subsequently sequenced on an Illuminia NextSeq 550 instrument (1 x 50 bp).

Following sequencing, library barcode splitting, read collapse, and trimming of adaptor sequences was performed using scripts from the icSHAPE pipeline (https://github.com/qczhang/icSHAPE) (40). Trimmed reads were first mapped to a costume genome that includes rRNA and repeated genomic sequences. Reads that did not map to the costume genome were then aligned to the human genome (hg38) using STAR (v 2.7.6a) (35, 36). We identified peaks using MACS2 (41). Peaks were required to appear in at least 2 replicates for further analysis. We extended the MACS2 identified peak summits to 100 nt to simulate the IP peak and performed intersection with the hg38 longest transcript isoforms using bedtools (v 2.30.0) (42). We identified peak changes using a generalized linear model and the DESeq2 (38) and QNB (43) programs in R (v 3.6.3). To control for changes in gene expression, gene level expression changes were subtracted from changes in IP peak reads for significantly modified peaks (44). Motif enrichment was calculated using HOMER(45) while metagene plots were constructed using MetaPlotR (46). GO-term annotation and enrichment was performed using Metascape (47) and GO-terms were collapsed with REVIGO (48) and plotted in R.

### SLAM-seq

Metabolic labeling to determine RNA half-life followed the method of Herzog *et al*. (49) UOK262 cells were seeded the day before the experiment at a density that would allow for exponential growth for the duration of the experiment (approx. 50% confluency). The morning of the experiment, UOK262 media was removed and replaced with UOK262 media supplemented with 100 µM 4-thiouridine (4SU; Carbosynth #NT06186) to begin the “pulse”. 4SU-containing media was exchanged every 8 hr during the 24 hr “pulse” period to increase the efficiency of labeling. Following 24 hr labeling, the “pulse” media was removed. Cells were washed 2X with PBS. Cell culture growth medium containing 10 mM uridine (i.e., 100X excess over 4SU; Sigma Aldrich #U3003-5G) was added to the cells to begin the “chase”. At select time points of interest, “chase” media was removed, and cells were lysed in using Trizol Reagent (ThermoFisher #15596026) according to the manufacture’s protocol with the following modifications: a) RNA isolation was performed in the dark and b) DTT (Sigma Aldrich #43816) was added to the aqueous phase (final concentration 0.1 mM) to keep samples under reducing conditions. RNA iodoacetylation was performed according to Herzog *et al.* To each reaction, we added 6 µg of RNA, 5 µL of 100 mM iodoacetamide (SigmaAldrich #I6125), 5 µL of 500 mM NaPO_4_, pH 8.0, 25 µL of DMSO and nuclease-free water to a final volume of 50 µL. Reactions were incubated in the dark for 15 min at 50°C. Reactions were quenched with the addition of 1 µL of 1 M DTT. RNA was isolated by ethanol precipitation and resuspended in a final volume of 10 µL and RNA quality control was performed on RNA High Sensitivity TapeStation (Agilent).

Library preparation was performed using QuantSeq 3’-mRNA library prep kit for Illumina (Lexogen #015) using the manufacturer’s protocol and recommended maximum input amount (500 ng). Library sequencing was performed on a HiSeq 3000 machine in PE100 mode. Following the recommendation of Herzog *et al.,* read 2 was discarded. Demultiplexed fastq files were applied to the SLAMDUNK pipeline (https://github.com/t-neumann/slamdunk) (Herzog *et al.* v 0.4.2) (49) and mapped against hg38. The optional --multimappers flag was enabled. Following the pipeline, the tcount output file was collapsed used the included SLAMDUNK Alleyoop collapse module. Curve fitting was performed according to a first-order decay reaction in R (v 3.6.3).

### Ribosome Profiling

Protocols were adapted from Arango *et al.* (50) and Ingolia *et al*. (51). Cells were plated in triplicate at a seeding density of 1.5 million cells per plate on 15-cm plates and grown to 80-90% confluency. At collection, media was aspirated, and cells were immediately flash frozen by submerging the bottoms of dishes in liquid nitrogen for 5-10 sec. Cells were maintained on dry ice until collection. Cells were collected by scraping on wet ice using 500 µL Lysis buffer (10 mM Tris-HCl, pH 7.4, 5 mM MgCl_2_, 100 mM KCl, 1% Triton X-100, 2 mM DTT, and 100 μg/mL cycloheximide (Millapore-Sigma #C4859-1ML)). Lysis buffer was transferred from one plate to the next to collect the cells from all three plates (approx. 1.5 total volume). Cell lysates were then triturated 10 times with a 25-gauge needle and clarified by centrifugation at 20, 000 x *g* for 10 min at 4°C. Lysates were aliquoted in 500 µl and stored at −80°C until use.

Each lysate was treated with 0.3 U/μL (1.5 µL for 500 µl of lysate) of RNase I (ThermoFisher # AM2294) for 40 min at 20°C, while the remaining lysate was flash frozen in liquid nitrogen for later total RNA isolation. Following the RNase I digestion, 7.14 μL of SUPERase-In (ThermoFisher # AM2694) were added to each sample, and the samples were loaded onto 15%–45% sucrose gradients that were made in 5 mM Tris-HCl pH 7.4, 37.5 mM KCl, 0.75 mM MgCl2 buffer and by using a BioComp Gradient Master (BioComp), and centrifuged at 37, 000 rpm in 4°C for 2 hr. Samples were then fractionated with BioComp fractionator. Fractions containing 80S monosomes were precipitated with 1 mL of 100% ice-cold ethanol at −80°C. After precipitation, monosome containing fractions were pooled and resuspended in 250 µL of Trizol Reagent (ThermoFisher #15596026). Sample workup was performed with the Direct-Zol kit (Zymo Research #R2050) following the manufacturer’s protocol. Samples were dried by vacuum freezing and quantified for ribodepletion.

Input samples were prepared from a 200 µL aliquot of the starting lysates in 800 µL of Trizol Reagent according to the manufacture’s protocol. Following phase separation, the aqueous phase was applied to a RNeasy Mini Kit (Qiagen # 74106) and RNA was purified according to kit instructions. Eluted input samples were then DNase treated using 4U of Turbo DNaseI (ThermoFisher #AM2239) in 100 µL total reaction volume for 30 min at 37°C. Following DNase treatment, reactions were cleaned on a RNeasy Mini Kit (Qiagen # 74106) following kit instructions. Samples were dried by vacuum freezing and quantified for ribodepletion.

For ribodepletion, 10 µg of each replicate (in 2 x 5 µg reactions) was ribodepleted using the RiboPool ribosome RNA *Homo sapiens* depletion kit (siTools Biotech #054). We supplemented this pool using custom-made oligos (designed by siTools) designed to be specific for our UOK cell line (1 µL of standard kit oligo + 1 µL of 1:30 dilution of custom oligos). The manufacturer’s instructions were followed for ribodepletion. Duplicate reactions were pooled at the final elution step. Following ribodepletion, ribosome protected fragments (RPFs) were cleaned using a Zymo RNA cleanup kit (Zymo Research #R1016) following the manufacturer’s protocol. The final elution was performed in 2 x 6 µL nuclease-free water. Inputs were spiked with 2 µL of 1:10 diluted ERCC Spike-in mix (ThermoFisher #4456740) and cleaned using a Zymo RNA cleanup kit (Zymo Research #R1016) following the manufacturer’s protocol. The final elution was performed in 2 x 6 µL nuclease-free water. The cleaned input reactions were subsequently fragmented using Ambion RNA Fragmentation reagent (ThermoFisher #AM8740) for 4 min at 95°C followed again by clean-up using a Zymo RNA cleanup kit and elution in 2 x 6 µL of water.

RPFs and inputs were electrophoresed on denaturing 15% Urea PAGE gels and RNA between 20 and 34 nt was size-selected. RNA was recovered from the gel by crush and soak and EtOH precipitation. RNA 3′-ends were dephosphorylated in a 10 µL reaction with 2 µL T4 PNK (NEB #M0201S) 1 µL of FastAP Thermosensitive Alkaline Phosphatase (ThermoFisher #EF0651), and 1 µL of RiboLock (ThermoFisher #EO0381) for 60 min in dephosphorylation buffer (70 mM Tris pH 6.5, 10 mM MgCl_2_, 1 mM DTT) with shaking. After 60 minutes, the components of the 3′-adaptor reaction were added (1 µL pre-adenylated and 3′-adaptor, 1 µL of 10X RNA ligase Buffer, 1 µL of T4 RNA Ligase (NEB #M0437M), 1 µL of 100 mM DTT, and 6 µL of PEG 8000) and the reaction was incubated overnight with shaking at 16°C.

Ligations were cleaned with a Zymo RNA cleanup kit (Zymo Research #R1016) and excess adaptor was removed through electrophoresis on a 15% denaturing Urea-PAGE gel. Following gel purification, RNA was reverse transcribed with Superscript IV reverse transcriptase (ThermoFisher #18090050) according to the manufacturer’s protocol. Following RT, samples were pooled as follows: WT RPFs, Mutant RPFs, WT inputs, and Mutant inputs. Pooled samples were cleaned using the Zymo DNA Clean and Concentrator kit (Zymo Research #D4014) according to the manufacturer’s protocol. Cleaned pools of samples were gel purified to remove excess primer and eluted cDNA was cleaned through Zymo DNA Clean and Concentrator kit (Zymo Research #D4014).

The cDNA from twice-cleaned pooled samples was circularized in at 15 µL reaction containing 12 µL cDNA, 0.5 µL circLigase II (Lucigen # CL9025K) 1.5 µL 10X circLigase buffer, 0.75 µL of 50 mM MnCl2 and nuclease-free water to 15 µL. Reactions were incubated for 2 hr at 60°C and purified by Ampure XP beads (Beckman #A63881). Libraries were PCR amplified in a 30 µL reaction containing 14 µL circularized library, 15µL Phusion HF 2X master mix (NEB #M0531L), 0.75 µL of 20 µM P3/P6 tall primer mix (see supplemental table XX) and 0.25 µL of 25X Sybr Green (ThermoFisher #S7563). Libraries were amplified for 8-12 cycles as determined by RT-qPCR on a QuantStudio3 RT-qPCR thermal cycler. Amplified libraries were cleaned up using Ampure XP beads and eluted in 10 µL of nuclease-free water. Three additional cycles of PCR were performed in 20 µL total volume containing 10 uL Phusion HF 2X master mix and 250 nM of P3/P6 Solexa primer mix and were again cleaned using Ampure XP beads. Samples were subject to a final gel purification through a 10% non-denaturing TBE PAGE, to maximize the purification of RPF and input libraries, and to remove empty adapter bands. Samples were submitted for sequencing on NextSeq 2000 P2, two independent times to obtain enough reads for analysis. All primers are listed in Table S1.

Following sequencing of the input and IP samples, barcode splitting, read collapse, and trimming of adaptor sequences was performed using scripts from the icSHAPE pipeline (https://github.com/qczhang/icSHAPE) (40). Trimmed reads were first mapped to a costume genome that includes rRNA and repeated genomic sequences. Reads that did not map to the costume genome were then aligned to hg38 using STAR (v 2.7.6a) (35, 36). Mapped reads were counted by htseq-count program (v 0.9.1) (37) and differential analysis was performed using DEseq2 (38) in R (v 3.6.3).

### miRNA-seq

Total RNA was extracted from cells using Trizol Reagent (ThermoFisher #15596026) according to the manufacture’s protocol. Following phase separation, the aqueous phase free water. Decapped RNA was subsequently digested to nucleosides (53) in a 30 µL reaction containing 24 µL RNA, 1 µL of 2U/µL Nuclease P1 (Sigma Aldrich, #N8630), 3 µL 1M ammonium acetate, pH 5.2, and 100 fmol of ^13^C-adenosine (Cambridge Isotope #CLM-3678-0.05) as an internal standard. Reactions were digested for 3 hr at 45°C. After 3 hours, 3µL of 1M ammonium bicarbonate and 1 µL of 0.002 U/µL of snake venom phosphodiesterase from *Crotalus ademanteus* (Sigma Aldrich #P3243) were added to the reaction and incubation continued for 2 hr at 37°C. After 2 hr, 1 µL of 1U/µL bacterial alkaline phosphatase (ThermoFisher #18011015) was added to the reaction and the reaction continued for 1 hr at 37°C. Digested nucleosides were passed through a 3 kDa MWCO spin filter (Amicon # UFC500396) and the filter was rinsed with 3 x 200 µL nuclease free water before being dried under vacuum. Dried samples were dissolved in 45 µL Buffer A (0.1% formic acid). Nucleosides were separated on a Luna Omega C18 reverse-phase column (1.6 um PS, 100 Å, 30 mm, ID 2.1 mm, Phenomenex) with a flow rate of 0.200 µL/min under the following conditions. Buffer A = 0.1% formic acid in HPLC-grade water; Buffer B = 0.1% formic acid in acetonitrile with the gradient described in Table S3. Following chromatography, nucleosides were resolved on a SciEX 500 QTOF in positive electrospray ionization mode using a Multiple-reaction-monitoring (MRM) MS method over the range of 100-350 daltons. (Spray voltage = 5500 V; Curtain gas = 35; CAD gas = 7; Temp = 600 °C; Declustering potential = 60 V; Collision energy = 10 V; accumulation time = 0.1 sec) The nucleosides were quantified using the nucleoside to base ion mass transitions and analyzed as the ratio of m^6^A was applied to a RNeasy Mini Kit (Qiagen) and RNA was / A or m^6^A / A. The precursor and product ions are listed in purified according to kit instructions. Libraries were prepared using the QIAseq microRNA Library Kit (Qiagen #331502) and sequenced using Illumina NovaSeq SP run with 100 base single-end reads. Reads for each sample were trimmed for adapters and low-quality bases using cutadapt (v 3.4) (34) and umitools (v 1.1.1) (52). Reads were mapped to hg38 using STAR (v 2.7.6a) (35, 36) and were counted by htseq-count (v 0.9.1) (37). Differential analysis was performed using DEseq2(38) in R (v 3.6.3).

### RNA Mass Spectrometry

Total RNA was extracted from cells using Trizol Reagent (ThermoFisher #15596026) according to the manufacture’s protocol. Following phase separation, the aqueous phase was applied to a RNeasy Mini Kit (Qiagen) and RNA was purified according to kit instructions. Total RNA underwent 2 rounds of polyA selection using NEB oligo-d(T)_25_ beads and protocol, followed by 1 round of ribodepletion (ThermoFisher human/mouse transcriptome isolation kit #K155002) following the manufacturer’s protocol for incubation and ethanol precipitation. RNA was decapped for 4 hr at 37°C in a total volume of 50 µL containing 500 ng RNA, 5 µL 10X NEB Buffer #2 (NEB #B7002S), 5 µL RppH (NEB #M0356S). Decapped RNA was cleaned using Zymo Oligo clean and concentrator kit (Zymo Research #D0460) and manufacturer’s protocol. The final elution was performed in 2 x 12 µL of nuclease-Table S4.

### Metabolite Mass Spectrometry

Cellular metabolites were extracted in 80 % ice-cold methanol. The TCA metabolites were derivatized with 3-Nitrophenylhydrazine (3NPH; Sigma Aldrich #N21804). In a 500 µL plastic vial, 15 µL each of the extract (or standard), internal standard (IS), 1-ethyl-3-(3-dimethylaminopropyl) carbodiimide (EDC; Sigma Aldrich #39391), and 3NPH solutions were mixed. After heating at 40°C for 30 min, the vials were cooled on ice and 7 µL of butylated hydroxytoluene (BHT; Sigma Aldrich #PHR1117) was added. After centrifugation, the supernatant was transferred to injection vials. Liquid chromatography (LC) were performed with a Shimadzu 20AC-XR system. MS/MS was performed with a TSQ Quantiva triple quadrupole mass spectrometer (Thermo Fisher Scientific) operating in SRM mode with negative ionization. The injection volume was 3 µl. Separation was achieved at 60°C with a 2.1 x 100 mm, 2.7 µm Cortex C18 column (Waters). Mobile phase A was 0.01% formic acid in water and mobile phase B was 0.01% formic acid in acetonitrile. The flow rate was 300 µL/min and the metabolites were separated with a gradient (0-0.2min/5%B; 12min/60%B; 12.1-13min/95%B; 13.1-15min/5%B). The TCA metabolites and amino acids in the cell extracts were determined by calibration curves generated by the Thermo Xcalibur software. Calibration curves, constructed by plotting the peak area ratios vs. standard concentrations, were fitted by linear regressions with 1/x weighting. The peak area ratios were calculated by dividing the peak areas of the target compounds by the peak areas of the corresponding isotopic internal standards.

### m^1^A primer extension

We adapted the protocol from Kawarada *et al.* (2017) (54). Briefly, we designed oligos for primer extension using published sequences (54) and included a 5’-conjugated infrared (IR) dye (IR700 for mt-tRNA-Arg and IR800 for mt-tRNA-Lys) on the oligos (IDT). Each reaction was carried out with 2 to 3 µg of total RNA. Total RNA was mixed with 50 fmol of oligo and 1 µl of buffer (50 mM Tris-HCl pH 8.0, 5 mM EDTA) in a 5 µl total reaction. Samples were heated to 80°C for 2 min, and then allowed to cool to room temperature at a slow ramp rate of 0.1°C/sec to allow RT oligos to anneal to target RNAs. While oligos annealed, ddNTP (Sigma Aldrich #GE27-2045-01) mixes were prepared. For mt-tRNA-Arg, 5 mM of ddATP was mixed with 5 mM of dTTP and dGTP, whereas for mt-tRNA-Lys, ddCTP was mixed with dTTP and dGTP. Once the RNA-oligo samples cooled, 1 µl of the corresponding d/ddNTP mix, 1.5 µL of 25 mM MgCl_2_, 0.5 µL of SuperScript III reverse transcriptase (ThermoFisher #18080044), and 2 µL of SuperScript III 5X first strand buffer was added to each reaction. Samples were incubated for 1 hr at 47°C. Samples were cleaned using Zymo Oligo Clean and Concentrator kit (Zymo Research #D4061) following the manufacturers protocol, eluted in 2 x 6 µl nuclease-free water. Samples were mixed with equal volume of 2X RNA loading buffer (ThermoFisher #R0641). Samples were denatured by heating for 1 min at 95°C and then electrophoresed on denaturing 20% urea page gels. Gels were visualized on an infrared detecting Licor scanner. All primers are listed in Table S1.

### m^1^A Misincorporation Analysis

Total RNA was extracted from cells using Trizol Reagent (ThermoFisher #15596018) according to the manufacture’s protocol. Following phase separation, the aqueous phase was applied to a RNeasy Mini Kit column (Qiagen #74106) and RNA was purified according to kit instructions. Following purification, the 28S and mt-ND5 transcripts were reverse transcribed 20 µL reaction using transcript-specific primers downstream of the m^1^A sites with Thermostable Group II Intron Reverse Transcriptase (TGIRT; InGex #TGIRT50) following the manufacturer’s protocol. Completed reactions were cleaned on a Zymo DNA Clean and Concentrator kit (Zymo Research #D4014). Transcripts were amplified with barcoded oligos (Table S1) targeting either 28S or mt-ND5 for 8-14 cycles as determined by RT-qPCR on a QuantStudio3 RT-qPCR thermal cycler. Reactions were purified by Ampure XP beads (Beckman #A63881) following the manufacturer’s protocol. DNA quality was assessed on Agilent D1000 tape station. Barcoded samples were pooled and sequencing on an Illuminia Micro Miseq in paired-end read mode. Demultiplexed reads were aligned to hg38 using STAR (v.2.7.6a) (35, 36). The presence of m^1^A was quantified using samtools-mpileup (55) to create individual base-pair frequencies for every position within the mt-ND5 and 28S rRNA transcripts. A window of 5 nucleotides centered on the m^1^A site was assessed. Misincorporation frequency was defined as the percentage of non-canonical base reads at a given site, with a higher misincorporation frequency corresponding to a higher frequency of modification. Misincorporation frequency was quantified by the sum of the frequencies of mis-read bases divided by the total read depth of a position.

### Transwell Migration Assay

The transwell assays were performed using an adapted version of the protocol described in Justus *et al.* (56). Cells were serum starving cells overnight using 1% FBS. The following day, 15, 000 cells were plated evenly across each Permeable Support for 24-well plate with 8.0mm Transparent PET Membrane (Corning #353097) in triplicate, using a total of 100 mL of media. Next, 600 mL of complete media (containing 10% FBS), serving as a chemoattractant, was added to the lower chamber, while 600 mL of low serum (1% FBS) media was added to the lower chamber of the control wells. After 24-48 hr of incubation in normal culture conditions, cells were removed from the upper chamber using a cotton tipped swab and fixed for 15 min in 70% ethanol. Cells were then stained for 30 min with 1% crystal violet aqueous solution (Sigma Aldrich #V5265) and washed several times in nuclease-free water. Membranes were allowed to dry before being imaged using the Lionheart FX Automated Microscope (Biotek). Migrated cells were then counted using ImageJ (v1.52k) and statistical analysis and plots were generated with R (v 3.6.3).

### Reagents and Biological Resources

For a complete list of Biological Resources and Software, please refer to Table S5.

### Statistical Analyses

All statistical tests, meaning, and values are listed in the main text or figure legends. The following statistical tests were used as appropriate: Student’s two-tailed t-test, Fisher’s exact test, Mann-Whitney U test, two-tailed binomial test, two-tailed Z test, ANOVA with multiple comparison assuming non-Gaussian distribution.

### Materials and Data Availability

The raw sequencing data obtained from the RNA-seq, miRNA-seq, m^6^A-IP, ribosome profiling, SLAM-seq and reported in this paper have been deposited in GEO (Accession number GSE228565). The high-throughput sequencing data from patient samples was previously generated and is available at GEO at GSE157256. High-throughput sequencing data was processed using freely available software. Scripts used to process data and generate plots can be found at <u>https://</u> <u>github.com/BatistaLab/UOK_manuscript</u>. For a complete list of software, please refer to Table S5.

## RESULTS

### Establishment of a system to study fumarate accumulation

Loss of FH activity and subsequent fumarate accumulation drive tumor development in HLRCC patients (15, 18). To determine the effect of fumarate accumulation on levels of RNA methylation, we used a lentiviral system to generate isogenic cell lines that are distinguished solely by FH activity. To achieve this, we expressed either a FLAG tagged wild-type (UOK262-WT) or inactive version of FH (UOK262-MUT) in UOK262 cells (UOK262-PAR), an immortalized renal tumor cell line derived from a HLRCC patient (15) (Figure 1A). The FH transgene expressed in UOK262-MUT cells carries the same FH Q396P mutation present as the parental UOK262 cells, which abrogates enzymatic activity leading to the accumulation of high levels of fumarate (15). Although the isogenic cell lines express FH protein at equivalent levels (Figure 1B top, Figure S1A), only the expression of wild-type FH restores enzymatic activity (Figure 1B middle, Figure S1B). In gel activity of the downstream TCA cycle enzyme, malate dehydrogenase, is similar between the two cell lines, demonstrating sample preparation preserved enzymatic activity (Figure 1B bottom and Figure S1C).

**Figure 1:**
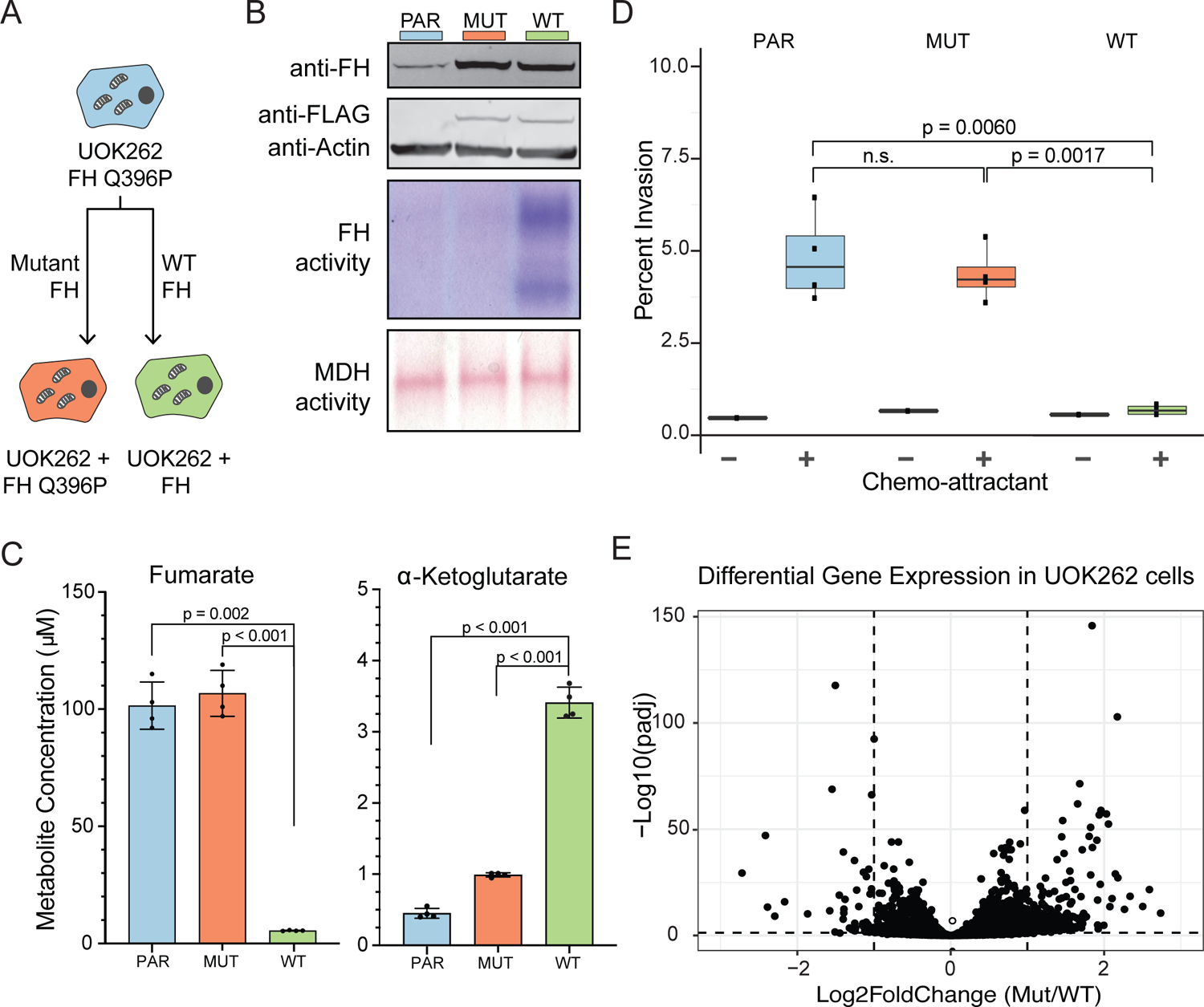
A cellular system to study the effect of fumarate accumulation on gene expression. (A) Cell lines used in the present study. The patient derived UOK262 cell line, which carries the fumarate hydratase (FH) Q396P mutation (blue) was used to generate isogenic cell lines that express either a WT (green) or mutant (orange) version of the FH protein. This color scheme is used throughout the manuscript. (B) Top Panel: Level of FH protein expressed in parental, UOK262-MUT, or UOK262-WT cells as determined by western blot. Actin is used as a loading control. Middle Panel: FH activity in UOK262 PARENT, MIT, and WT cell lines determined by an MTT-based colorimetric in-gel activity assay. Bottom Panel: Malate dehydrogenase (MDH) activity of UOK262 parental, mutant, and WT cell lines determined by an MTT-based colorimetric in-gel activity assay. MDH activity was used as a loading control for FH activity. (C) LC-MS/MS analysis of fumarate (left) and αKG metabolite (right) levels from UOK262-PARENT, UOK262-MUT, and UOK262-WT cell lines. Error bars indicate mean ± s.d. from n = 4 replicates and statistical significance was determined by student’s t-test. Values to generate this plot are included in Table S6. (D) Analysis of invasion potential of UOK262-PARENT, UOK262-MUT, and UOK262-WT cell lines was measured on a transwell invasion assay. Error bars indicate mean ± s.d. for n=3 replicates and statistical significance was determined by t-test. (E) Volcano plot depicting steady state fold change in mRNA levels between UOK262-MUT and UOK262-WT cells. Changes in gene expression are determined with DESeq2. Values used to generate this plot are included in Table S8.

Expression of the wild-type FH protein results in a 50-fold reduction in the level of fumarate and a significant increase in αKG compared to the UOK262-PAR and UOK262-MUT cell lines (Figure 1C). Previous work has demonstrated that a high αKG/succinate ratio promotes the activity of DNA and histone demethylases, and that modifying this ratio is sufficient to regulate multiple chromatin modifications (20, 57). In our isogenic system the ratio between αKG and succinate changes approximately 30-fold from 0.143 in UOK262-MUT cells to 4.4 in UOK262-WT cells (Figure S1D, Table S6). The ratio observed in UOK262-WT cells is comparable to the ratio measured in the non-cancerous kidney cell lines HK2 (ratio = 2.0) and HEK293 (ratio = 5.1) (Table S6). Together, this suggests that the metabolic environment in the UOK262-MUT cells disrupts the activity of 2OGDDs which may contribute to changes in downstream modifications.

We examined whether the wild-type transgene restored the defected TCA cycle in the UOK262 parent cells. FH functions in the TCA cycle by catalyzing the interconversion of fumarate to malate. Metabolite levels in the TCA cycle (Figure S1D), including malate, are restored by expression of wild-type FH enzyme (Figure S1D). Additionally, in FH-deficient cells, argininosuccinate is generated from fumarate via reversal of the Urea cycle enzyme argininosuccinate lyase (58). As expected, argininosuccinate levels in the UOK262-MUT cells were elevate compared to the UOK262-WT cells (Figure S1E) (58, 59).

Restoration of the TCA cycle also results in increased in the levels of ATP in UOK-262-WT cells (Figure S1F). Disruption of the TCA cycle often leads to high dependency on glucose for catabolism and anabolism (15) in cell lines. In agreement with previous research, the UOK262-PAR and UOK262-MUT cells are highly dependent on glucose for growth, while expression of wild-type FH in UOK262 decreases glucose dependency (Figure S1G). Accumulation of fumarate caused by FH mutations can enhance cellular migration and invasion properties (25, 60). Consistent with this observation, we found that restoration of FH activity significantly abrogates transwell cell invasion (Figure 1D) (61), suggesting that metabolic rewiring supports oncogenic phenotypes. Together, these results demonstrate that our isogenic system serves as a suitable model system to study the effects of fumarate accumulation on RNA modifications.

To further characterize these cells, we performed gene expression profiling. Although introduction of the flag-tagged FH Q396P mutation has minimal impact on the transcriptome of the UOK262-PAR cells (Figure S1H, Table S7), the presence of functional FH (UOK262-WT) results in significant changes in gene expression (Figure 1E). Overall, we observe 1359 genes with significant changes in gene expression (padj < 0.05) between UOK262-MUT, and UOK262-WT cells (Table S8). Among the differentially expressed genes, we identified transcripts coding for proteins involved in chemotaxis, regulation of cell migration, and cell-cell adhesion (Figure S2A). In agreement with previous work, we found genes in the PI3K-Akt signaling and the NRF2 pathways, which are dysregulated in UOK262 cells in response to fumarate accumulation (61, 62), to be enriched among genes differentially expressed between UOK262-MUT and UOK262-WT cells (Figure S2B, Figure S2C). Previous work identified overexpression of microRNA hs-miR-200b-3p as driver of EMT in cells that accumulate fumarate (25). In our isogenic cell system, we observed modest differences in miRNA levels between UOK262-MUT and UOK262-WT cells (Figure S2D) (Table S9). Additionally, we found no significant differences in mRNA level for hs-miR-200b-3p or hs-miR-200b-3p targets (Figure S2E)

### Expression of 2OGDD RNA demethylases is not affected by metabolic rewiring

A high αKG-to-succinate ratio is necessary for the activity of αKG-dependent dioxygenases, and changes in this ratio have been shown to affect gene expression (20, 57). Competitive inhibition of 2OGDDs by fumarate has been proposed to drive cellular transformation by inhibiting the activity of enzymes in this family. Five enzymes in this family act upon methylated RNAs, raising the possibility that RNA methylation levels can change in response to fumarate accumulation. An *in vitro* study found that the affinity of 2OGDD family members to fumarate, succinate and 2HG differs between individual enzymes (63). Therefore, we rationalized that the effect of fumarate accumulation on different RNA modifications might be varied. Of the 2OGDD enzymes that target RNA ((Figure 2A), we evaluated the expression level of ALKBH1, ALKBH5, and FTO, which have been previously linked to cancer. When comparing UOK262-WT and UOK262-MUT cells, we observed no differences at either the mRNA or protein level of ALKBH1, ALKBH5 or FTO (Figure 2B, S2F, S2G). Similarly, the expression level of ALKBH1, ALKBH5 or FTO mRNAs is not significantly different when comparing normal renal cortex, primary and metastatic HLRCC sites in patient data (Figure 2C, Figure S2H, Table S10, Table S11) (39). Thus, metabolic rewiring does not seem to impact the expression level of the ALKBH family members ALKBH1, ALKBH5 or FTO.

**Figure 2:**
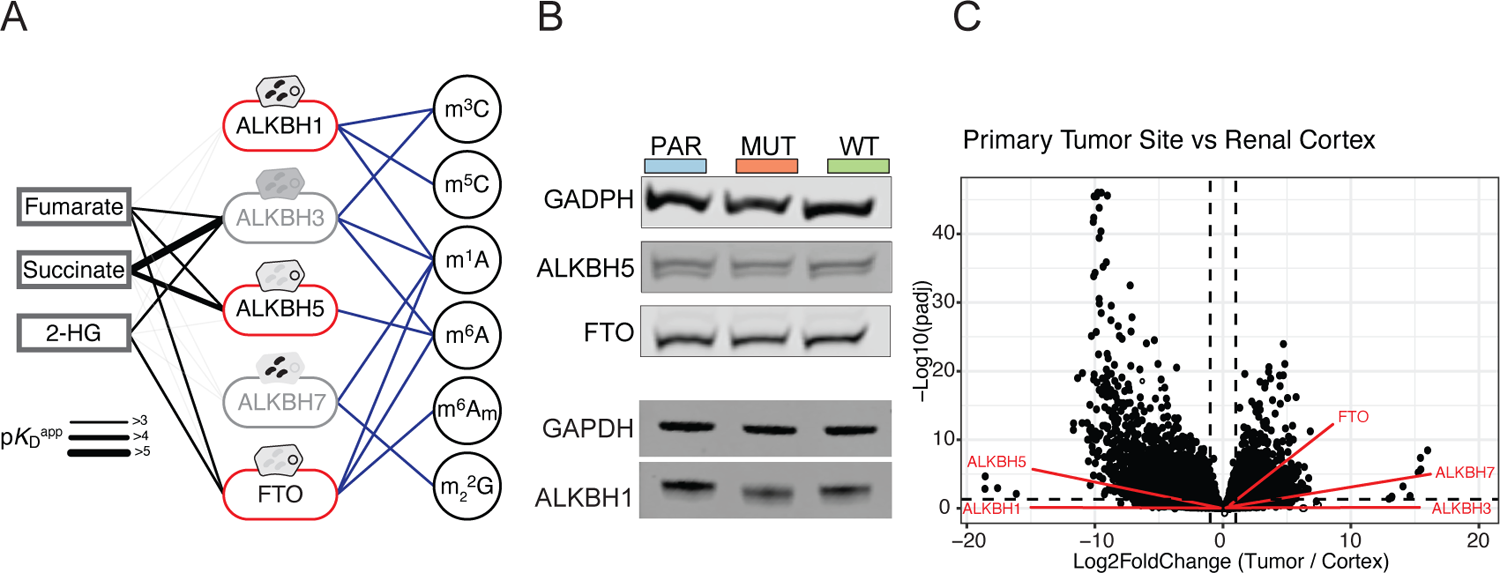
Changes in gene expression in response to fumarate accumulation. (A) Scheme depicting the 2OGDD family members that are known to act on RNA modifications. Those that are the subject of this study are highlighted. (B) Representative western blot of demethylases ALKBH1, ALKBH5, and FTO in UOK262 parental, mutant, and WT cell lines. GAPDH is used as a loading control. (C) Volcano plot of gene expression changes in patient mRNA samples between primary tumor and renal cortex sites. Changes in gene expression are determined with DESeq2. ALKB demethylases are highlighted in red. Values used to generate this plot are included in Table S10.

### ALKBH1 target modifications do not respond to fumarate accumulation

ALKBH1 is known to localize to the mitochondria, where fumarate levels are expected to be the highest (64–66). ALKBH1 removes m^1^A from mt-tRNAs and plays a critical role in mitochondrial respiration by regulating the conversion of 5-methylcytosine (m^5^C) to f^5^C in the anticodon of mitochondrial tRNA-Methionine (mt-tRNA-Met) (Figure 3A, S3A) (54, 67). As ALKBH1 is exposed to high levels of fumarate, we examined whether the levels of mitochondrial substrates of ALKBH1 increased in response to fumarate accumulation. Using a recently reported f^5^C sequencing assay to assess the presence of f^5^C in mt-tRNA-Met (68), we observe no differences in f^5^C levels between UOK262-MUT and UOK262-WT cells (Figure 3B). This implies that f^5^C levels do not respond to fumarate accumulation and suggests that ALKBH1 is not sensitive fumarate. Silencing of ALKBH1 expression confirmed the dependency of the f^5^C signal on ALKBH1 activity (Figure 3B, 3C, 3D). In addition to f^5^C, ALKBH1 demethylates the m^1^A modification in mt-tRNA (Figure S3A). Upon silencing ALKBH1 in our isogenic system, we did not observe changes in m^1^A levels on mt-of fumarate or succinate inhibits 2OGDD enzymes found in the cytoplasm and nucleus (8, 22–25, 70–74). We sought to determine if loss of FH activity and subsequent accumulation of fumarate inhibits the activity of 2OGDDs that target RNA and localize outside of the mitochondria, such as ALKBH5 and FTO (Figure 4A). In this case, inhibition of these enzymes would result in mRNA m^6^A hypermethylation. Indeed, by LC-MS/MS we observe that high levels of fumarate correlate with increased levels of m^6^A in the mRNA fraction of the UOK262-MUT cells (Figure 4B). In addition to m^6^A, FTO can also remove N6, 2′-O-dimethyladenosine (m^6^A). However, by LC-MS/MS we did not observe a statistically significant difference between UOK262-WT and UOK262-Mutant m^6^A tRNA^Arg^ or mt-tRNA^Lys^ (Figure S3B, S3C), precluding us from determining if fumarate accumulation impacts levels of m^1^A in a ALKBH1 dependent manner. Taken together, these data indicate that ALKBH1 activity is unaffected by metabolic changes accompanying the loss of FH.

**Figure 3:**
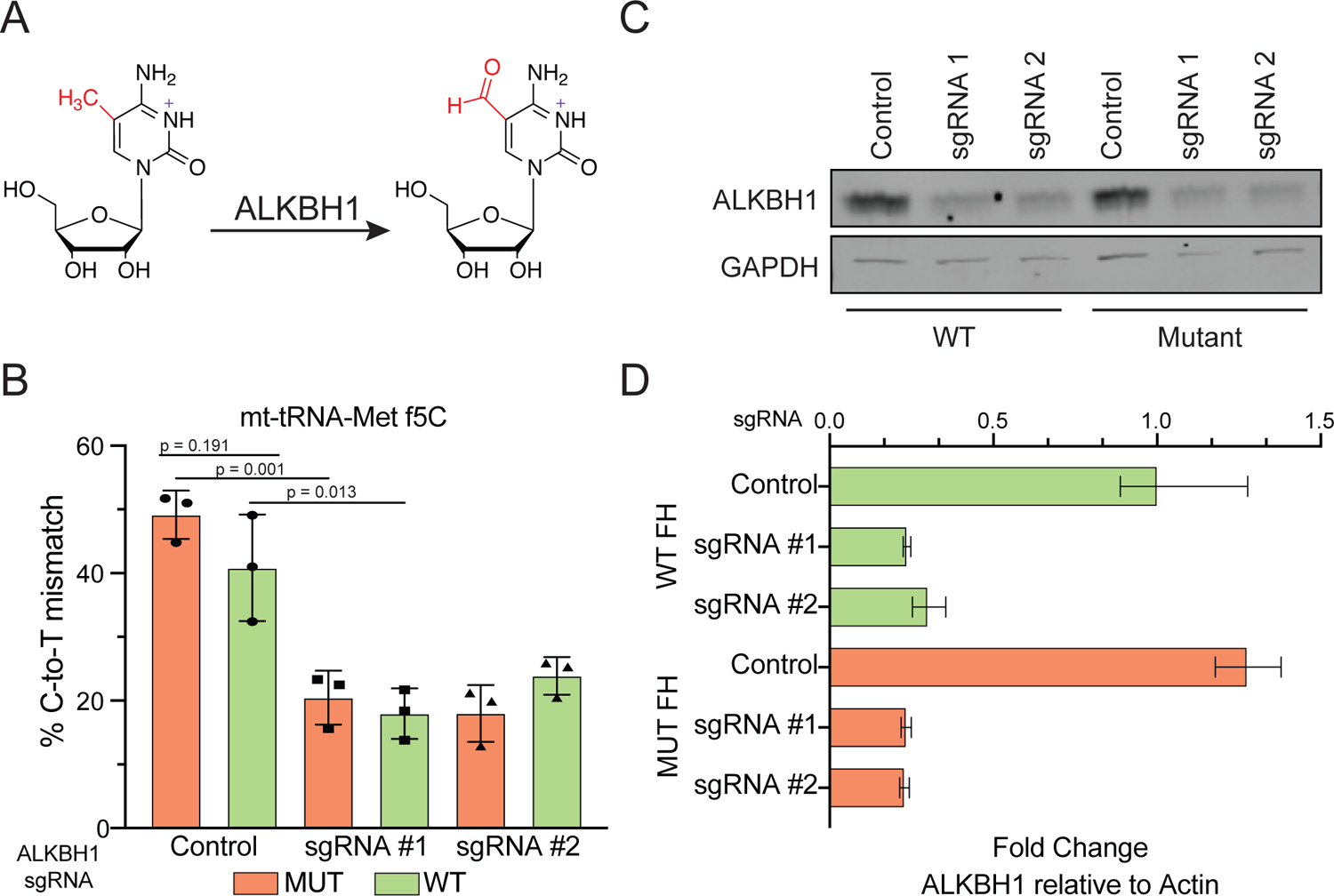
Changes in ALKBH1-dependent demethylation in response to fumarate accumulation. (A) Scheme depicting the oxidation of 5-methylcytosine to 5-formylcytosine by ALKBH1. (B) Percentage of C to T mis-incorporation at position 34 of mt-tRNA-Met for 5-formylcytosine in UOK262-MUT (orange) and UOK262-WT (green) cells. Misincorporation was measured in control cells containing non-targetting sgRNAs, as well as two independent ALKBH1 KD cell lines. (C) Representative western blot of ALKBH1 KD in UOK262-MUT and UOK262-WT cell lines. ALKBH1 was independently knocked down with two sgRNAs. GAPDH is used as a loading control. (D) RT-qPCR of ALKBH1 KD in UOK262-MUT (orange) and UOK262-WT (green) cells. Error bars indicate mean ± SD for n = 3 replicates.

**Figure 4:**
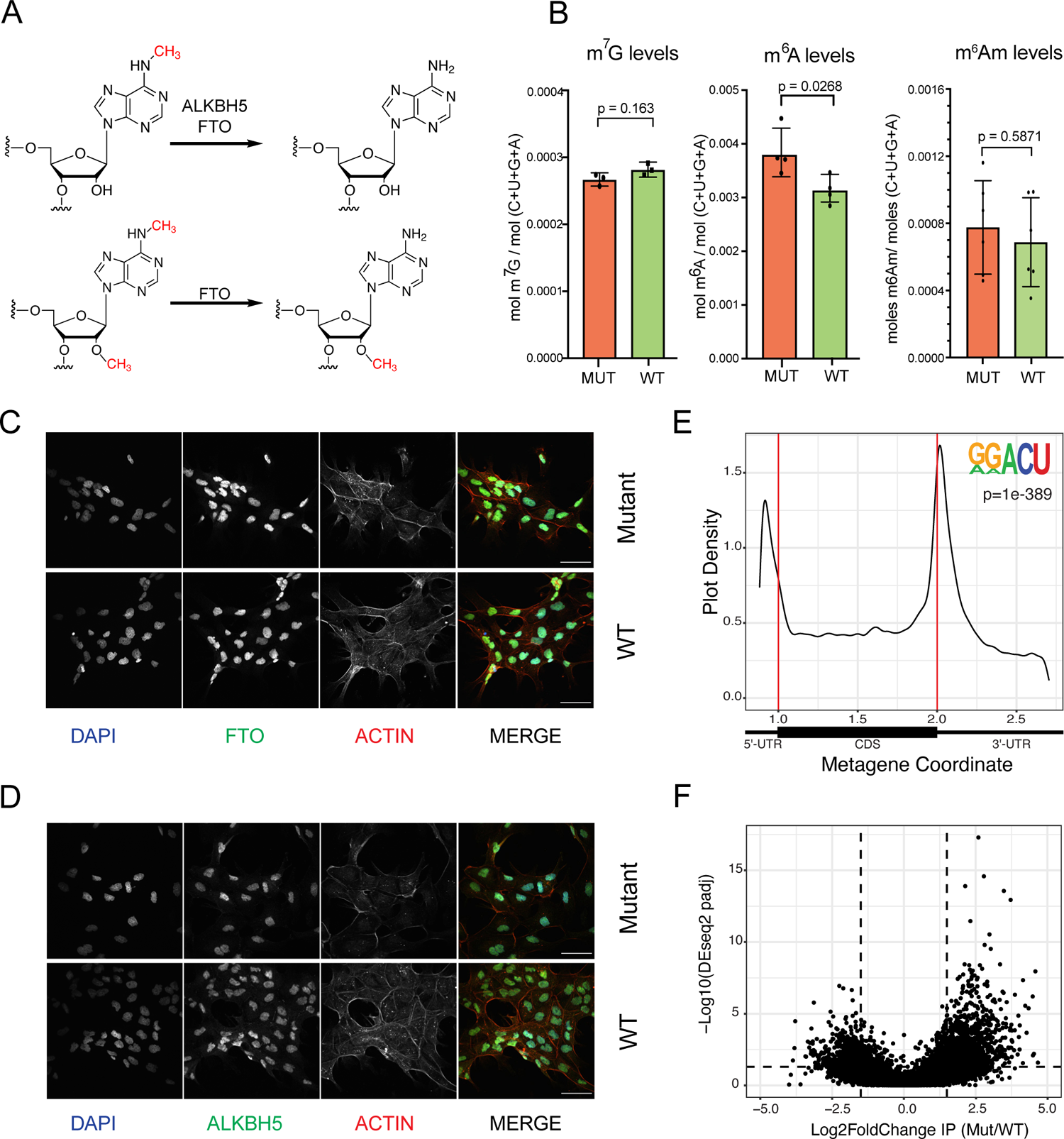
Changes in m6A and m6A-demethylase expression and localization in response to fumarate accumulation. (A) Scheme depicting the demethylation reactions performed by ALKBH family demethylases FTO and ALBKH5. (B) LC-MS/MS quantification of m^7^G, m^6^A, and m^6^A in polyadenylated RNA in UOK262-MUT and UOK262-WT cells. Error bars indicate mean ± SD for n = 4 replicates and p-values were determined by student’s t-test. (C) Representative immunofluorescent images of subcellular localization of FTO (green) in UOK262-MUT and UOK262-WT cells. The nucleus was stained with DAPI (blue) and cytoplasmic localization was visualized by actin (red). Scale bars = 10 µM. (D) Representative immunofluorescent pictures of subcellular localization of ALKBH5 (green) in UOK262-MUT and UOK262-WT cells. The nucleus was stained with DAPI (blue) and cytoplasmic localization was visualized by actin (red). Scale bars = 10 µM. (E) Metagene plot of peaks identified across transcripts in UOK262-MUT and UOK262-WT cells and motif enrichment analysis showing the most significantly enriched sequence across peaks identified with MACS2. *De novo* motif enrichment was performed with HOMER, p = 1e-120). (F) Volcano plots depicting fold change of reads in m^6^A peaks (m^6^A differential modification) in UOK262-MUT to UOK262-WT cells. Changes in m^6^A expression are present in Table S12.

Finally, we wondered whether fumarate accumulation affects mitochondrial mRNA methylation. Several transcripts, including mt-ND5, are known to contain the m^1^A modification (69). We leveraged the property of m^1^A to induce nucleotide misincorporation during cDNA synthesis to determine if m^1^A at mt-ND5 responds to fumarate. Targeted sequencing of the mt-ND5 m^1^A site showed no differences in cDNA misincorporation between UOK-MUT and UOK-WT, although overall the levels of misincorporation were relatively low (Figure S3D). Sequencing of an amplicon containing position 1302 in 28S rRNA, which is known to contain the m^1^A modification at high stoichiometry, showed high level of misincorporation in both UOK262-MUT and UOK262-WT cells (Figure S3E), confirming the assay can detect m^1^A misincorporation.

### Fumarate inhibits mRNA demethylation

While fumarate is thought to accumulate to highest levels in the mitochondria, it has been shown that accumulation levels (Figure 4B). Considering the levels of FTO andALKBH5 are similar between UOK262-MUT and UOK262-WT cells, one explanation for the elevated m^6^A levels could be due to changes in expression of the m^6^A methyltransferase complex. However, we observe no differences in mRNA or protein expression of the core components of the m^6^A methyltransferase complex (METTL3 and METTL14) when comparing UOK262-MUT and UOK262-WT cells (Figure S4A, S4B), supporting the hypothesis that changes observed in m^6^A levels result from inhibition of demethylase activity by fumarate. While we do not observe changes in the protein levels of the core components of the methyltransferase complex or the demethylases (Figure 2A, S4A), differences in m^6^A levels could be derived from changes in demethylase localization, as demethylase localization has been shown to impact substrate choice and activity (75). Through a combination of cellular fractionation and immunohistochemistry we determined that loss of FH activity does not affect localization of FTO or ALKBH5. In both UOK262-MUT, and UOK262-WT cells FTO is found mainly in the nucleus, where the preferred substrate is m^6^A (75), while ALKBH5 is found in both nucleus and cytoplasm (Figure 4C, 4D; S4C). The same localization pattern for FTO and ALKBH5 can be observed in human renal cortex and UOK262 xenograft sections (Figure S4D).

Together, these results show that accumulation of fumarate in HLRCC inhibits the m^6^A erasers FTO and ALKBH5 and leads to RNA hypermethylation.

### M6A-IP identifies changes in m^6^A sites

To identify sites of RNA hypermethylation dependent on fumarate accumulation, we performed m^6^A-immunoprecipitation (m^6^A-IP-seq) on both UOK262-MUT and UOK262-WT. In both cell types, the known m^6^A motif (RRACH) was identified through *de novo* motif discovery in regions enriched after m^6^A immunoprecipitation (Figure 4E). In agreement with previous studies, we observed a higher density of m^6^A enriched regions near the stop codon and the beginning of the 3’UTR (Figure 4E, Figure S4E) (76, 77). The peaks (across 6956 genes) in cells with active FH (UOK262-WT) (Collectively, we refer to the base modifications detected through m^6^A-IP as m^6^A(m), although most are likely m^6^A). When we performed differential m^6^A methylation analysis comparing UOK262-MUT and UOK262-WT peaks (44, 81), we identified 1129 peaks with higher m^6^A(m) signal (padj < 0.05; log2FoldChange (l2fC) >= 1.5) in cells that accumulate fumarate (UOK262-MUT) as compared to UOK262-WT(Figure 4F). These peaks are distributed across 943 transcripts, and the majority of these m^6^A peaks (800 peaks) are detected exclusively in UOK262-MUT, while the remaining 143 peaks are detected in both cell types. The number of upregulated peaks per transcript ranges from 1 to 5, with the majority (806 transcripts) having a single upregulated peak (Figure S4F). The m^6^A antibody can also capture the m^6^A modification, which Additionally, we identified a small subset of transcripts (192) occurs exclusively in adenosines next to the mRNA cap (78–80). We observe an enrichment of m^6^A signal near the TSS, which could reflect sites of m^6^A modification (Figure 4E). We identified a total of 15501 m^6^A(m) peaks (across 7248 genes) in FH-deficient cells (UOK^MUT^) and 15122 m^6^A(m) where all detected m^6^A peaks are upregulated. Of these, 13 transcripts have multiple m^6^A peaks, while on the remaining transcripts we detected a single m^6^A peaks. Conversely, we identified 499 peaks with lower m^6^A levels (padj < 0.05; l2fC <= −1.5) in cells that accumulate fumarate (Figure 4F). Of the 443 genes with peaks with lower m^6^A levels, 56 transcripts have m^6^A sites where all peaks are downregulated.

Given the size of the RNA fragments in our IP experiment, peaks within the first 100 nt of the transcript could represent regulation of cellular death and differentiation (Figure S5A, S5B, S5C and S5D, Table S14). Comparison of mRNA half-life between wildtype and mutant cells shows a trend of shorter half-life for transcripts in cells that accumulate fumarate (mean half-life Mutant = 5.69 hr; WT = 6.14 hr, KS-sites of m^6^A modification, which can also be recognized by test p = 5.13e-11) (Figure 5C). While we observe decreased the m^6^A-antibody. In our data sets, approximately 20% (3672) of peaks are found within the first 100 nt of the transcript (Figure S4G)). Of the 3553 transcripts with a peak in the first 100 nt, about 46% (1631 transcripts) have previously been mRNA stability in UOK262-MUT cells, these differences are not specific to transcripts with fumarate dependent changes in m^6^A levels (Figure 5C, LS-means = 0.441). Thus, we conclude that fumarate-dependent gain of m^6^A is not driving identified as m^6^A modified (Table S12) (82–84). Among the loss of mRNA stability in UOK262-MUT cells. 1129 upregulated peaks, 137 peaks (55 previously identified as m^6^A modified) are found in the first 100 nucleotides and Finally, to determine if fumarate dependent changes in are potential m^6^A sites. Finally, for the transcripts identified m^6^A levels affect translation, we examined ribosome with a single upregulated site, 53 are potential m^6^A sites. Overall, the m^6^A-IP-seq data is in agreement with the observed increase of m^6^A on a global level and identifies transcripts with increased levels of m^6^A modification.

**Figure 5:**
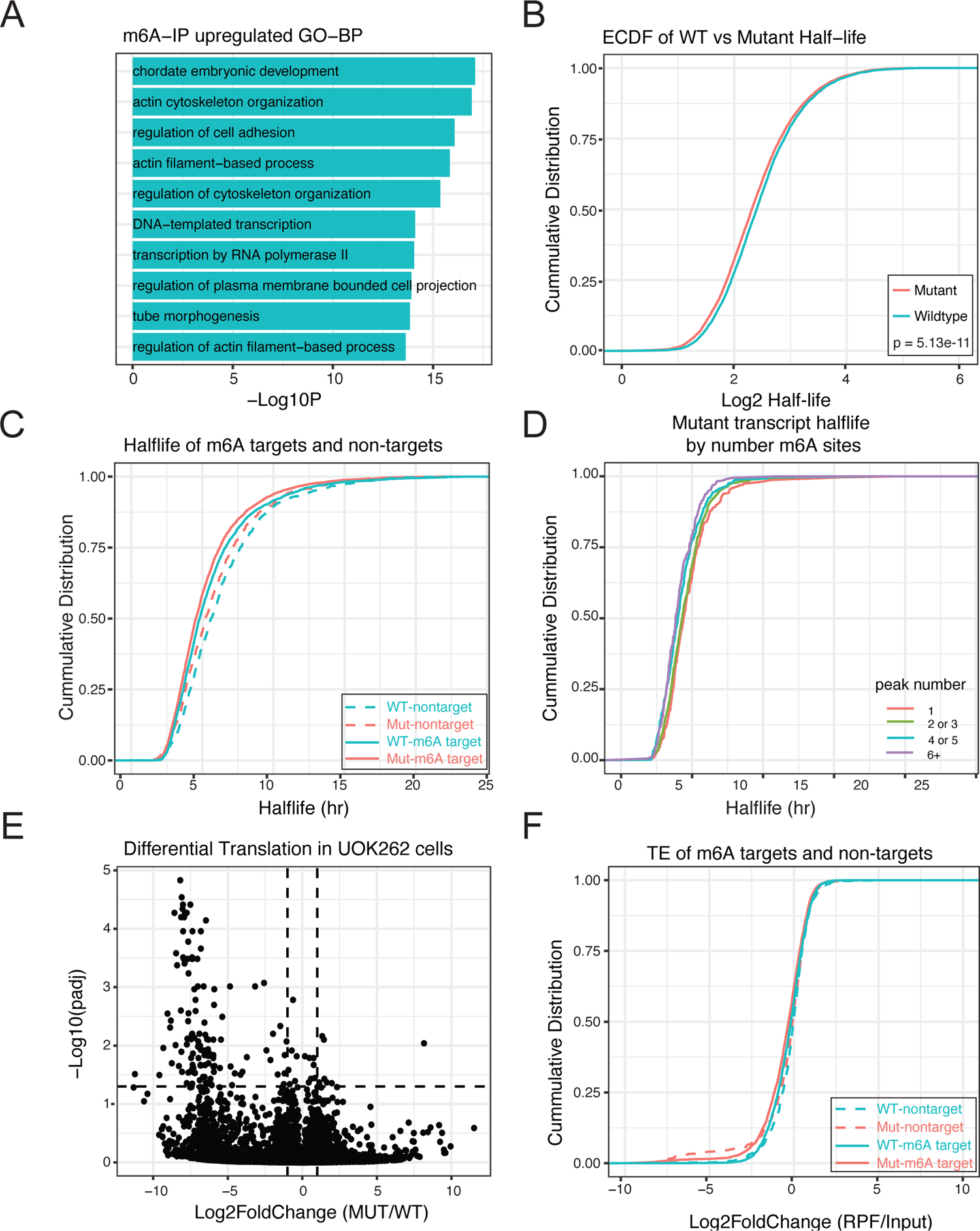
Transcriptome-wide mapping of m^6^A shows changes in cells with fumarate accumulation. (A) GO-biological term enrichment analysis for genes that are upregulated (l2fc > 1.5 and padj < 0.05; blue) in m^6^A-IP data. Depicted are the top 10 enriched GO-terms. (B) Cumulative distribution function (CDF) of high-confidence transcript stabilities for approx. 6, 800 transcripts (n=3). P-value was determined with KS-test. (C)CDF plot depicting mRNA half-life of m^6^A-target and non-target transcripts between UOK262-MUT and UOK262-WT cells. P-values were determined with KS-test. (D) Representative CDF plot depicting the half-life of transcripts from UOK262-MUT cells as a function of the number of m^6^A-peaks on each transcript. (E) Volcano plot depicting fold changes in mRNA translation between UOK262-MUT and UOK262-WT cells. Changes in ribosome occupancy for each transcript, relative to input levels, are determined with DEseq2. (F) CDF depicting Translational Efficiency of m^6^A-target and non-target transcripts between UOK262-MUT and UOK262-WT cells. P-values were determined with K.S. test. Values from half-life figures are available in Table S14 and values from riboseq figures are available in Table S15.

### Impact of fumarate dependent m^6^A hypermethylation on RNA metabolism

Transcripts with higher m^6^A(m) levels in the mutant are enriched for the biological process GO-terms cell adhesion (GO:0030155), and actin cytoskeleton organization (GO:0030036) (Figure 5A). Among the peaks with higher m^6^A signal we identified the gene Oxoglutarate Dehydrogenase (OGDH), a known target of ALKBH5. Impairment of ALKBH5 activity as a response to viral infection has been shown to lead to an increase of m^6^A modification on the transcript for OGDH (85). Additionally, since a fumarate-dependent invasion phenotype is one of the hallmarks of HLRCC, we asked if genes involved in Epithelial to Mesenchymal Transition (EMT), known to be an important driver of this phenotype, are represented among the hypermethylated genes. We identified 89 “EMT signature” genes among the transcripts with higher m^6^A levels in cells that accumulate fumarate (Table S13). Among these we identified vimentin, ZEB1, and ZEB2, genes previously implicated in EMT in UOK262 cells (25).

Previous literature implicates a role for m^6^A in mRNA stability and translation (reviewed in (86)). To understand if fumarate dependent changes in m^6^A levels affect mRNA stability in FH deficient cells we used metabolic labeling to determine transcript half-life in both UOK262-WT and UOK262-MUT cells. (Figure 5B, Table S14) (49). As previously described, we observed that m^6^A modified transcripts have a shorter half-life than transcripts with no m^6^A modification in both UOK262-WT (transcripts with no m^6^A modification vs modified transcripts: KS-test = p = 4.22 x 10^-15^) and UOK262-MUT cells (transcripts with no m^6^A modification vs modified transcripts: KS-test p = 2.2 x 10^-16^) (Figure 5C). Furthermore, transcripts with a higher number of m^6^A sites display a shorter half-life (Figure 5D). As observed in other studies (49, 87), transcripts with longer half-lives are enriched for genes associated with cellular macromolecule metabolism, while transcripts with shorter half-lives are enriched for genes associated with occupancy across the transcriptome through sequencing of ribosome protected fragments (Ribo-seq) (Table S15) (51). Comparing the translation efficiency (TE) in UOK262-WT and UOK262-MUT cells (Figure 5E), we observe minor differences between the two cell types. The majority of these differentially translated transcripts are found at low levels in the ribosome fraction. Finally, when we assessed the role of m^6^A modification in TE, we do not observe differences between m^6^A modified transcripts and transcripts with no m^6^A modification (Figure 5F). Together, these observations suggest fumarate accumulation does not substantially affect translation of m^6^A-containing RNAs.

## DISCUSSION

Throughout development, cells utilize distinct metabolic strategies to support growth, produce energy, and meet the unique demands of each tissue to maintain homeostasis (88, 89). In cancer, metabolic reprograming facilitates cellular proliferation and acquisition of new properties (3, 4). One instance of cancer-associated metabolic reprograming with a significant role in cancer is the shift to aerobic glycolysis driven by mutation of metabolic enzymes, such as the TCA cycle enzyme FH. This change results in altered cellular metabolite levels, which affect cellular signaling and gene expression (21). The shift in metabolism driven by loss of FH activity generates an inhibitory environment for enzymes in the 2OGDD family. This family of enzymes catalyzes hydroxylation reactions on proteins, nucleic acids, and lipids, and is involved in multiple pathways critical to maintaining cellular homeostasis (19). While previous studies have demonstrated inhibition of 2OGDDs acting on DNA, histones, and the HIF transcription factor by fumarate (8, 22–25, 70–74), no study has addressed the role of fumarate accumulation on RNA post-transcriptional modifications. Here, we sought to determine how the intracellular metabolic environment in HLRCC tumors, caused by loss of FH, affects the activity of ALKBH1, ALKBH5 and FTO, three RNA demethylases in the 2OGDD family. While these enzymes share a requirement for αKG, oxygen, and Fe^2+^ for catalysis, they have diverse substrate specificities and act on different classes of RNA transcripts (Figure 2A). As such, accumulation of fumarate can have a wide impact on RNA methylation.

### Accumulation of fumarate does not affect ALKBH1 activity

ALKBH1 is localized within the nucleus and mitochondria where it regulates the level of methyl modified nucleotides on cytoplasmic and mitochondrial tRNAs (54). In addition, ALKBH1 is required for the formation of f^5^C at position 34 homogenous across the transcriptome, as we observe a gain in m^6^A in less than 10% of total sites detected in our experiment. This specificity likely reflects the differences in affinity of FTO and ALKBH5 for each site across the transcriptome (75, 79, 95, 96). In contrast to m^6^A, we did not observe a statistically significant difference in global of the cytoplasmatic tRNA^Leu^ (CAA) and the mitochondrial m^6^A levels between UOK262-WT and UOK262-Mutant cells tRNA^Met^. On mt-tRNA^Met^ the cytosine at position 34 is methylated by NSUN3 to m^5^C and then converted to f^5^C by ALKBH1. Modification of the anti-codon loop of mt-tRNA^Met^ with f^5^C is required for translation of non-universal initiation codons (AUA and AUU) in mammalian mitochondria (54, 67, 90, 91). Loss of either NSUN3 or ALKBH1, and subsequent loss of f^5^C negatively affects mitochondrial function and has been linked to a number of diseases, including cancer (54, 67, 90, 92). Given the critical importance of ALKBH1 to mitochondria function, and the previous observations that mitochondrial function in HLRCC is impaired (15, 25, 60, 64), we hypothesized that accumulation of fumarate contributes to loss of mitochondria function through disruption of f^5^C levels. An analysis of f^5^C levels at the anti-codon loop of mt-tRNA^Met^ revealed no loss of f^5^C in cells that accumulate high levels of fumarate (Figure 3B). We propose that ALKBH1 is not sensitive to fumarate as an inhibitor, perhaps reflecting selective pressure on ALKBH1 to be functional in a cellular compartment that experiences accumulation of different levels of fumarate. Our observations in the UOK262 cell line agree with previous studies that measured druggability of enzymes in the 2OGDD family (63) and found ALKBH1 to be less susceptible to inhibition by αKG related oncometabolites (63).

### Accumulation of fumarate leads to mRNA m^6^A hypermethylation

While fumarate is thought to accumulate to highest levels in the mitochondria, it has been shown that accumulation of fumarate or succinate inhibits enzymes of the 2OGDD family found in the cytoplasm and nucleus resulting in pseudohypoxia, by stabilizing the transcription factor HIF1α (22, 70, 71) and hypermethylation of DNA (8, 23–25, 72–74). We therefore asked if FTO and ALKBH5, two enzymes of the 2OGDD family that localize to the nucleus or cytoplasm and act on mRNA, are affected by accumulation of fumarate. Both FTO and ALKBH5 are capable of demethylating m^6^A at internal positions (75), while FTO can additionally remove the m^6^A methyl group from m^6^A, a modification present in the extended cap structure of mRNA (82, 83). In accordance with potential inhibition of FTO, ALKBH5, or both by fumarate, we observed an increase in global mRNA m^6^A levels when cells accumulate fumarate (Figure 4B). We observe no changes in expression, or localization, of the methyltransferases or demethylases in our isogenic cell lines that could explain the differences we observe in m^6^A levels. This is consistent with previous work that has demonstrated that the ability of FTO to demethylate single-stranded DNA can be inhibited by fumarate (93) and inhibition of FTO by 2HG leads to mRNA hypermethylation (94). This increase of m^6^A is not (Figure 4B).

Hypermethylated transcripts include genes involved in multiple cellular processes, including cell migration and EMT. To understand how fumarate dependent changes in m^6^A levels impact gene expression in fumarate deficient cells we compared RNA stability and translation in UOK262-MUT and UOK262-WT cells. In our isogenic system, we did not find significant changes in half-life for the m^6^A modified population of transcripts between the two cell lines (Figure 5C). Yet, the observation that m^6^A modified transcripts tend to have shorter half-life in both UOK262-MUT and UOK262-WT cells, and that this effect depends on the number of m^6^A regions (Figure 5D), supports a role for m^6^A in controlling transcript stability in our isogenic system. The lack of impact of fumarate dependent m^6^A hyper methylation might be related to the distribution of hyper-methylated sites. The distribution of hypermethylated regions (peaks) is heterogenous, and in most hypermethyated transcripts, there is a single upregulated peak. It is possible that upregulation of m^6^A on only a fraction of possible sites on a transcript is not enough to affect its half-life. Across the transcriptome, while we see differences in RNA stability among transcripts with one and six or more m^6^A regions, differences between groups that differ by one or two m^6^A regions are modest. In fact, a recent study (97) has suggested that m^6^A-dependent regulation of half-life is complex, as only small differences in RNA expression levels were observed for groups of transcripts with high levels of methylation. It is also possible that fumarate dependent changes in m^6^A can affect RNA half-life, but not under the conditions used to culture the cells for our assays. It is worth pointing out that under proliferation conditions, in rich media containing an excess of glucose, the two cell lines have similar growth kinetics. It is only under conditions that mimic invasion that cells lacking FH activity show a phenotype. Under different conditions, such as during the process of invasion which is modulated by fumarate, differences in signaling pathways and other cellular processes could sensitize pathways controlling mRNA stability or translation to respond to fumarate dependent changes in m^6^A. In fact, across several systems loss of methyltransferase complex is tolerated under some conditions, such as proliferation, but blocks other cellular processes such as differentiation (98–102). Future work will determine if manipulation of ALKBH5 or FTO activity affects the fumarate dependent invasion phenotype of these cells.

### Accumulation of fumarate has different impacts on ALKBH family members

RNA demethylases in the αKG-dependent dioxygenase family act on diverse RNA modifications. We hypothesized that inhibition of these enzymes by accumulation of fumarate, or other structurally related metabolites, can globally remodel the RNA methylome. Given the differences in the binding affinity for αKG, or related TCA metabolites, between different RNA demethylases in the 2OGDD family, the effect on each type of RNA methylation will be different. In fact, previous work on the druggability of 2OGDDs found that ALKBH1, found in the mitochondria, is highly specific for αKG, and less susceptible to inhibition by the oncometabolites 2GH, succinate, and fumarate (63). Our observations that ALKBH1 targets are not affected by fumarate accumulation agree with this early study (Figure 2A). One hypothesis is that the demethylases with essential roles in mitochondria function have evolved to perform their vital functions in an environment with fluctuating levels of co-factors and potential inhibitors. In contrast, ALKBH5 and FTO have some sensitivity to both fumarate and succinate, two metabolites that accumulate because of loss of FH activity. The affinity of FTO for fumarate was found to be similar to that of EGLN1 (also known as PHD2), an enzyme shown to respond to fumarate accumulation (63). It is interesting to note that ALKBH5 is predicted to be more sensitive to the effects of fumarate than FTO, and to therefore hypothesize that targets of ALKBH5 may be more affected by fumarate accumulation. It remains to be determined if both ALKBH5 and FTO are impacted by excess fumarate. Not only the two enzymes have different affinities to αKG and fumarate, differences in localization between the two proteins could result in different levels of inhibition. FTO, which is exclusively nuclear, and the nuclear pool of ALKBH5 might be inhibited to different levels than the cytoplasmic pool of ALKBH5. Previous work has demonstrated that chromatin-associated RNA (caRNAs), including enhancer RNAs and retrotransposon elements, and mitochondrial polycistronic transcripts are methylated and substrate for enzymes in the ALKB group. Further work will be necessary to determine if other types of RNA modifications, or modifications on other types of RNA transcripts, are affected by fumarate accumulation.

In summary, our results report on the link between loss of FH and regulation of RNA methylation through the demethylases ALKBH1, ALKBH5, and FTO. While our results focus on fumarate accumulation, it is interesting to note that the diversity of metabolic rewiring observed in tumors, including mutations in different enzymes of the TCA cycle, can affect RNA post-transcriptional modifications in distinct ways. In addition, underlying tissue-specific factors, and the sensitivity of these enzymes to different inhibitors, may lead to pro-tumorigenic or non-growth promoting environments, as well as opportunities for future therapeutic interventions. The distinct impact of each metabolite on different enzymes opens the door to targeted inhibition and modulation of specific modifications.

## AUTHOR CONTRIBUTIONS

**Conceptualization:** CMF, PJB **Methodology:** CMF, DRC, JML, WML, PJB **Validation:** CMF, MDM, JCL, DC, SSM, ACS **Formal analysis:** CMF, MDM, JCL, DC, SSM, ACS, STG, CL, LMJ, DRC, PJB **Investigation:** CMF, MDM, JCL, DC, SSM, ACS, STG, CL, LMJ, DRC **Resources:** LMJ, WML, PJB **Data Curation:** CMF, MDM, SSM, PJB **Writing - Original Draft:** CMF, PJB **Writing - Review & Editing:** CMF, MDM, SSM, DRC, JLM, PJB **Visualization:** CMF, MDM, JCL, DC, SSM, PJB **Supervision:** JLM, WML, PJB **Funding acquisition:** CMF, JLM, WML, PJB.

## ACKNOWLEDGEMENTS

We thank Jaye Gardiner, Susan Gottesman, Sezen Meydan Marks, Khoa Tran, Joana Vidigal and members of her lab, and Sandra Wolin for critical reading of the manuscript. The authors thank the Center for Cancer Research (CCR) Genomics Core in Bethesda, Maryland, as well as the CCR Sequencing Facility in Frederick, Maryland, for help with high-throughput sequencing. The authors also thank the CCR Microscopy core for their help with the IF experiments. This work utilized the computational resources of the NIH HPC Biowulf cluster (http://hpc.nih.gov). We appreciate the editorial assistance of George Leiman. The views expressed in this article are those of authors and may not reflect the official policy or position of the National Institute of Health, Department of the Army, Department of Defense. Mention of trade names, commercial products, or organizations does not imply endorsement by the U.S. Government.

---

**Supplemental Figure 1:**
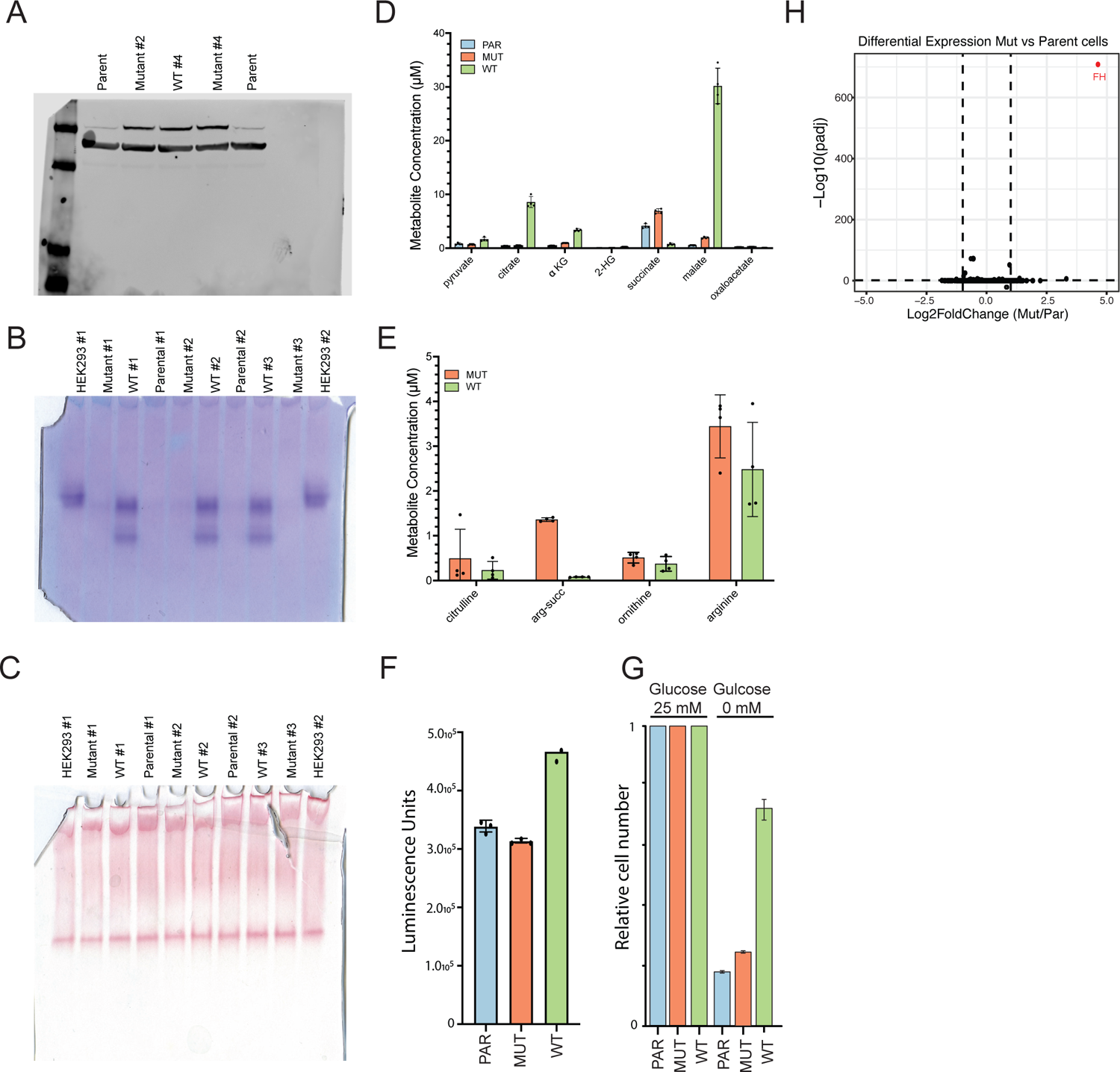
(A) Full size western blot of FH expression, as shown in Figure 1B. Actin (lower band) is used as a loading control. (B) Full size blot of FH colorimetric in-gel activity as shown in Figure 1B. Multiple replicates of Parental cells (negative control), HEK293 cells (positive control) and the UOK262 isogenic cell lines are present. (C) Full size blot of malate dehydrogenase colorimetric in-gel activity, as shown in Figure 1B. Multiple replicates of Parental, HEK293, and the isogenic cell lines are present. (D) LC-MS/MS quantitation of TCA cycle metabolites from UOK262 parental (blue), mutant (orange), and WT (green) cell lines. Error bars represent mean ± s.d. for n = 4 replicates and statistical significance was measured by t-test. Values used to generate this plot are in Table S6. (E) LC-MS/MS quantitation of Urea Cycle metabolites from UOK262 mutant (orange) and WT (green) cell lines. Error bars represent mean ± s.d. for n = 4 replicates and statistical significance was measured by students t-test. (F) ATP production in UOK262 parental (blue), mutant (orange), and WT (green) cell lines. Error bars represent mean ± s.d. for n = 3 replicates and statistical significance was measured by t-test. (WT vs Mut: p-value = 0.0018). (G) *In vitro* growth of UOK262 parental (blue), mutant (orange), and WT (green) cell lines in cell growth medium containing either 25 mM or 0 mM glucose. (H) Volcano plot depicting fold change in mRNA levels between UOK262-MUT and UOK262-PAR cells. Changes in gene expression are determined with DESeq2. The FH gene is highlighted in red. Values used to generate this plot are included in Table S7.

**Supplemental Figure 2:**
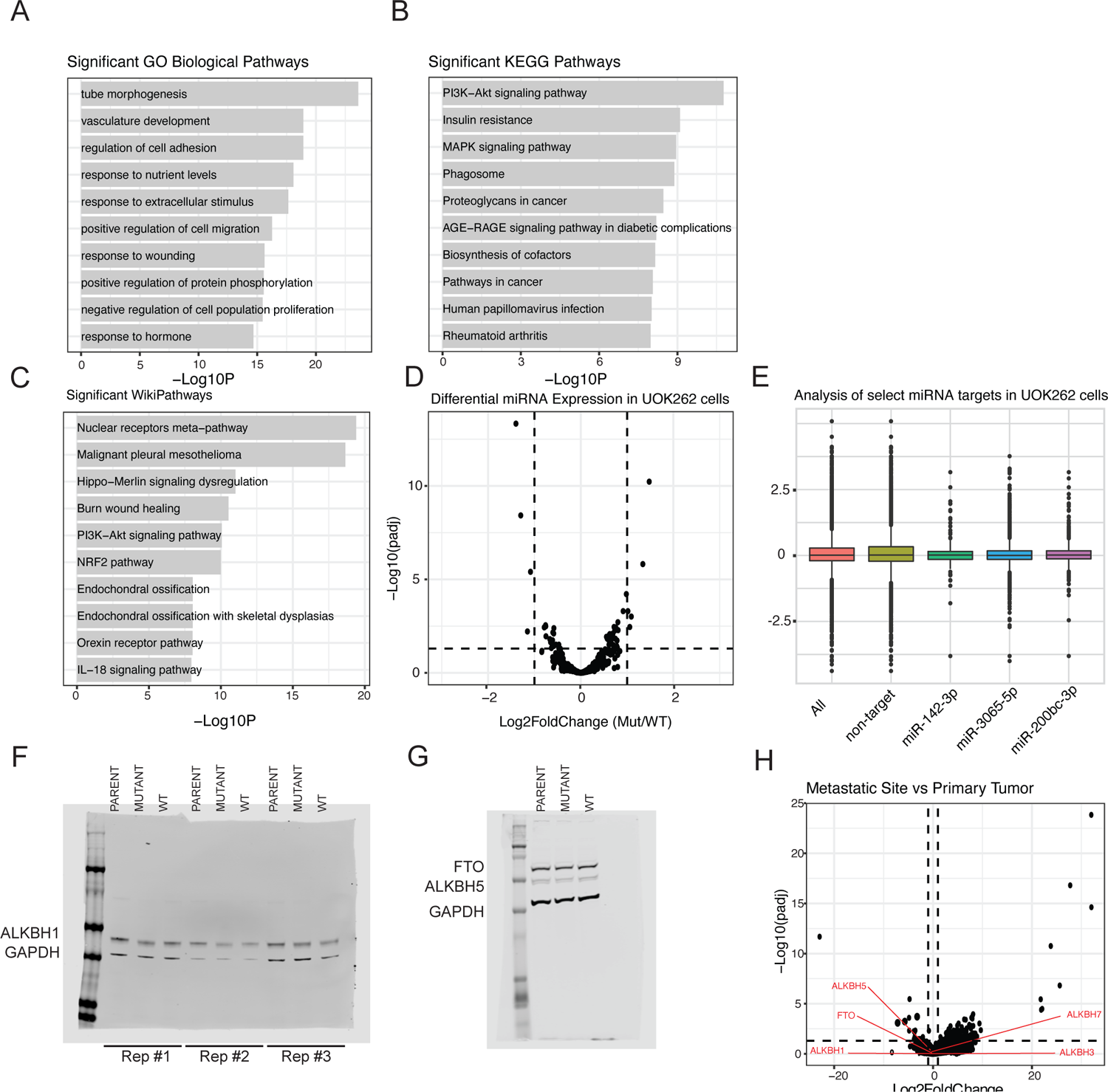
(A)GO-biological term enrichment analysis for genes that are upregulated (l2fc > 1.5 and padj < 0.05; blue) or downregulated (l2fc > 1.5 and padj < 0.05; red) in UOK262-MUT RNA-seq data. Depicted are those GO-terms with a log10 p-value > 5. (B) KEGG pathways enrichment analysis of genes that were downregulated (l2fc < −1.5 and padj < 0.05) in UOK262-MUT RNA-seq data. (C) KEGG pathway enrichment analysis of genes that were upregulated (l2fc > 1.5 and padj < 0.05) in UOK262-MUT RNA-seq data. (D) Volcano plot depicting fold change in miRNA levels between UOK262MUT and UOK262WT cells. Changes in miRNA expression are determined with DESeq2. Values used to generate this plot are included in Table S9. (E) Box plot depicting changes in miRNA targets of miR-142-3p (green), miR-3065-5p (blue) and miR-200bc-3p (violet) as compared to all genes (red) and non-target genes (yellow) in the UOK262-MUT vs UOK262-WT miRNA data. (F) Full size western blot for ALKBH1 in Figure 2B. (G) Full size western blot for FTO and ALBKH5 in Figure 2B. (H) Volcano plot of gene expression changes in patient mRNA samples between metastatic tumor and primary tumor sites. Changes in gene expression are determined with DESeq2. ALKB demethylases are highlighted in red. Values used to generate this plot are included in Table S11

**Supplemental Figure 3:**
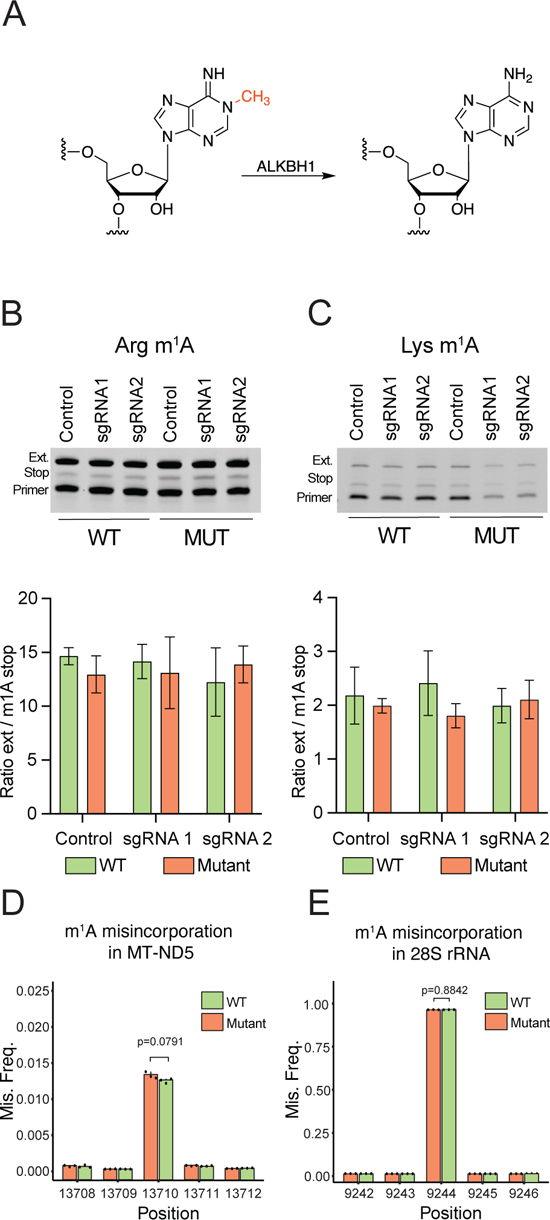
(A) Scheme depicting the demethylation of m^1^A to adenosine by ALKBH1. (B) Representative detection and quantitation of m^1^A from mt-tRNA-Arg at position A17 through primer extension. The primer is the lowest band, cDNA truncated at the m^1^A position appears in the middle, and the full cDNA product is the top band. Bands were quantified in ImageJ and error bars represent mean ± s.d. for n = 3 replicates. (C) Representative detection and quantitation of m^1^A from mt-tRNA-Lys at position A58 through primer extension. The primer is the lowest band, the extended cDNA for the detection of m^1^A appears in the middle, and the extension through the unmodified A is the top band. Bands were quantified in ImageJ and error bars represent mean ± s.d. for n = 3 replicates. (D) Misincorporation analysis of m^1^A site 9244 of 28S rRNA in UOK262-MUT (orange) and UOK262-WT (green) cells. Error bars indicate mean ± SD for n = 3 replicates and p-values were determined by student’s t-test. (E) Misincorporation analysis of m^1^A site 13710 of mt-ND5 in UOK262-MUT (orange) and UOK262-WT (green) cells. Error bars indicate mean ± SD for n = 3 replicates and p-values were determined by student’s t-test.

**Supplemental Figure 4:**
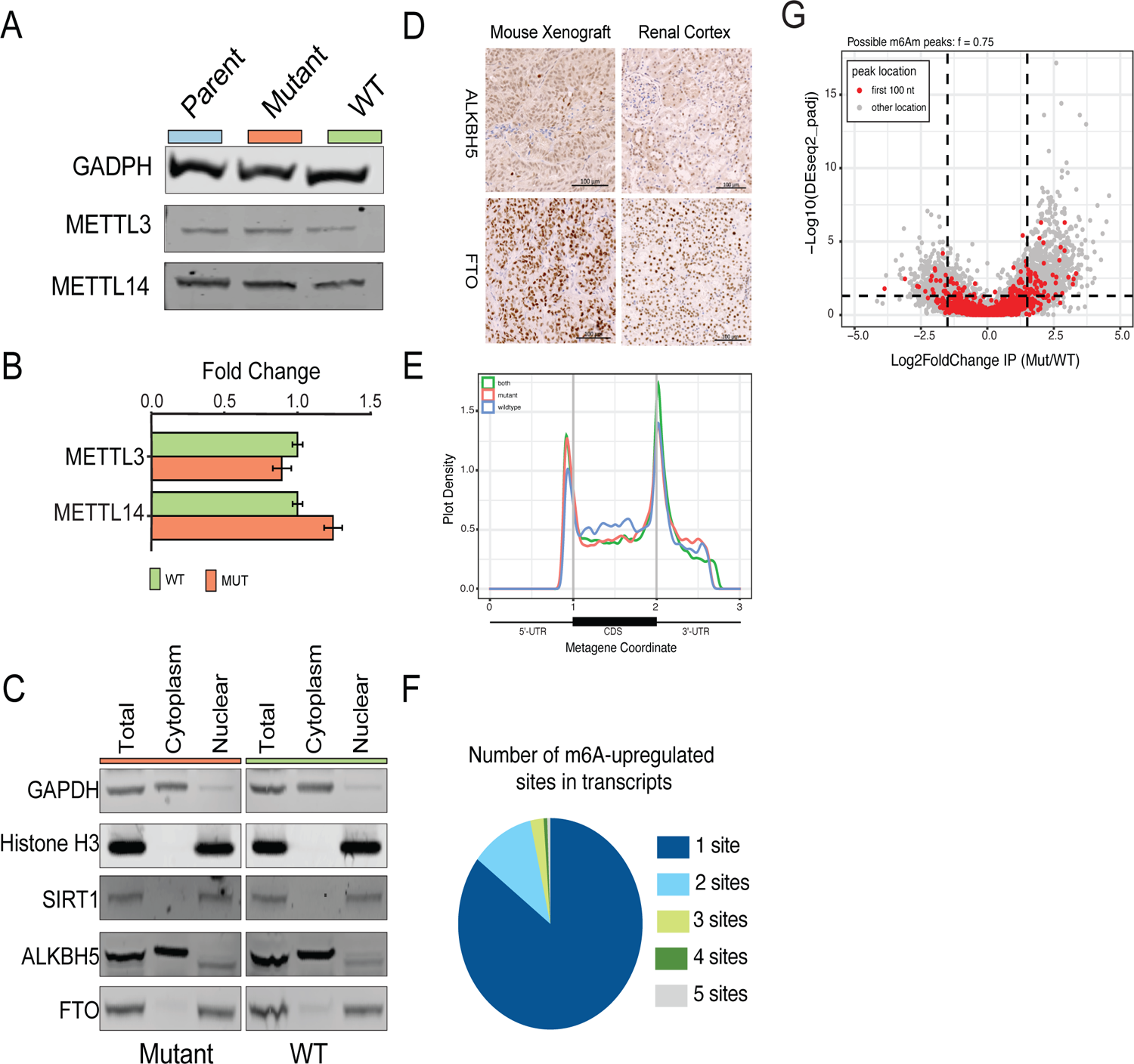
(A) Representative western blot of m^6^A writers in UOK262 PARENT, MUT, and WT cell lines. GAPDH is used as a loading control. (B) RT-qPCR of m^6^A writers and erasers in UOK262-MUT (orange) and UOK262-WT (green) cells. Error bars indicate mean ± s.d. for n = 3 replicates. (C) Western blot of sub-cellular fractionation of in UOK262-MUT and UOK262-WT cells. GADPH (cytoplasm), Histone H3 (nuclear pellet), and SIRT1 (nucleus) were used as controls. (D) Immunohistochemistry blot of UOK262 mouse xenograft models and human renal cortex of ALKBH5 (top) and FTO (bottom). Signal is detected in most cells in proximal and distal tubules. Glomerulus cells show both positive and negative cells. Scale bar = 100 µm. (E) Metagene profile of mRNAs displaying distribution of peaks that are unique to UOK262-WT (blue), unique to UOK262-MUT (red) or common to both cell lines (green). (F) Pie chart depicting the fraction of m^6^A sites per transcript. (G) Volcano plot depicting the log-fold change of possible m^6^A peaks. The peaks located within the first 100 nt of the transcripts are depicted in red. All other peaks are depicted in grey.

**Supplemental Figure 5:**
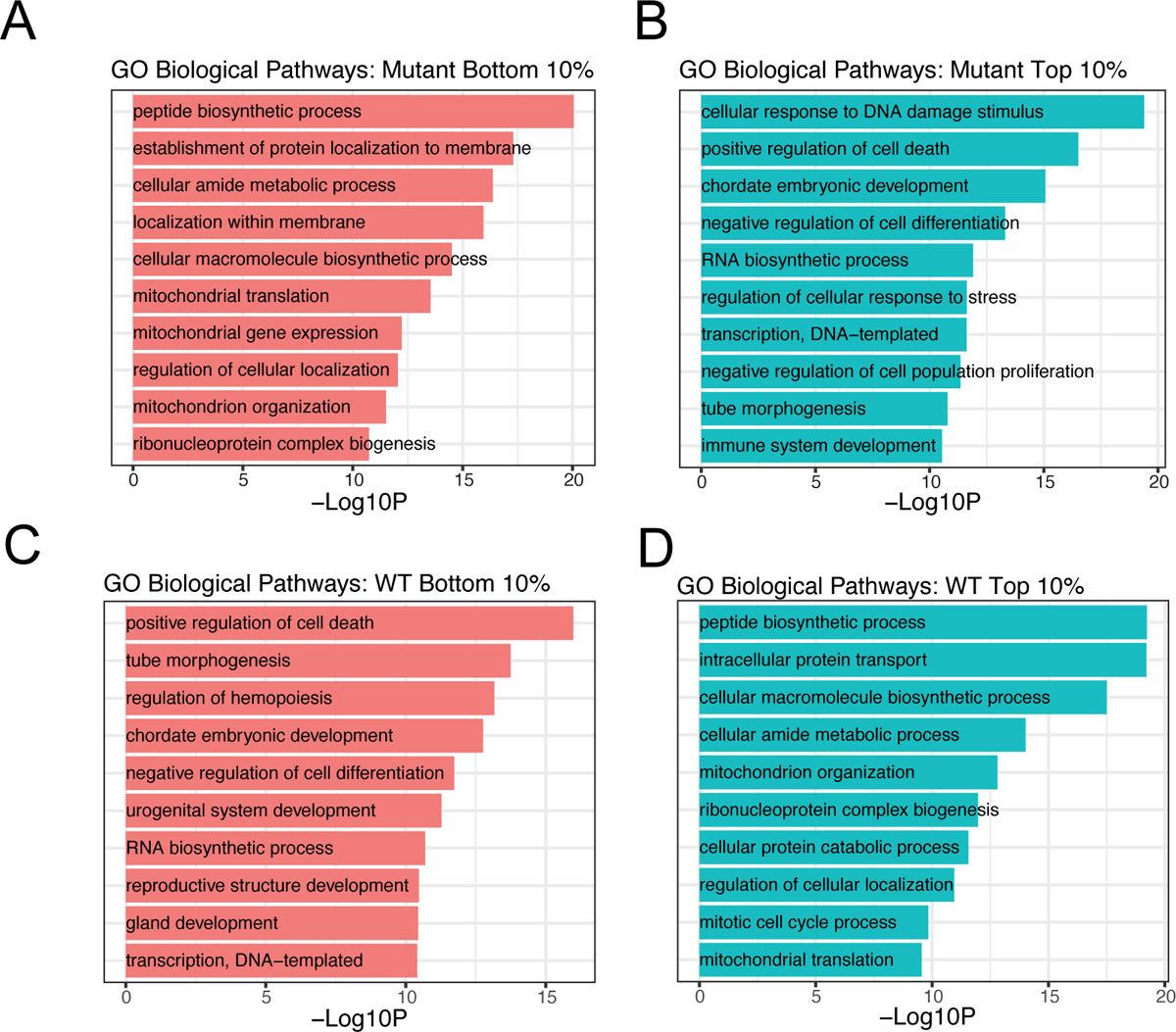
(A) GO-biological process analysis for genes that are in the bottom 10% for half-life in Mutant. Depicted are the top 10 GO-terms. (B) GO-biological process analysis for genes that are in the top10% for half-life in Mutant. Depicted are the top 10 GO-terms. (C) GO-biological process analysis for genes that are in the bottom 10% for half-life in WT. Depicted are the top 10 GO-terms. (D) GO-biological process analysis for genes that are in the top10% for half-life in WT. Depicted are the top 10 GO-terms.

## SUPPLEMENTAL TABLES

**Table S1:** List of oligonucleotides used in this study

**Table S2:** List of antibodies used in this study

**Table S3:** LC-MS/MS gradient parameters for RNA nucleotide analysis

**Table S4:** LC-MS/MS precursor and product ions for RNA nucleotides

**Table S5:** Complete list of biological resources and software in this study

**Table S6:** LC-MS/MS of TCA metabolites from UOK262, HEK293, and HK2 cells

**Table S7:** Differential expression analysis comparing gene expression changes between UOK262-MUT and UOK262-PAR cells.

**Table S8:** Differential expression analysis comparing gene expression changes between UOK262-MUT and UOK262-WT cells

**Table S9:** Differential expression analysis comparing miRNA expression changes between UOK262-MUT and UOK262-WT cells

**Table S10:** Differential expression analysis comparing gene expression changes between primary tumor and renal cortex sites

**Table S11:** Differential expression analysis comparing gene expression changes between metastatic and primary tumor sites

**Table S12:** Differential m^6^A expression between UOK262-MUT and UOK262-WT cells Differential m^6^A expression was calculated through read filtering and normalization of the peak-level expression changes to those of the gene-level input changes. Ensemble gene ID (ENSG_ID); common gene name (gene_name); numerical m^6^A peak ID calculated by MACS2 (peak_ID); chromosome location (chr); chromosome coordinates for m6A peak start (start); chromosome coordinates for m^6^A peak end (end); location of the peak on the forward or reverse strand (strand); calculated log2foldchange and padj value of the gene (gene_l2fc and gene_padj); calculated log2foldchange of the m6A peak (peak_ l2fc); peak_l2fc minus input gene_l2fc (diff_l2fc); padj values (DEseq_padj and logDEseqP); direction and significance of the padj and l2fc values (direction); presence and identity of the known RRACH motif for m^6^A modifications (m^6^A motif); is the gene known to be a target of m^6^A (target of m^6^A); calculated halflife of transcript in UOK262-MUT or UOK262-WT cells (half_Mutant and half_WT; see Table S14); calculated l2fc and padj from riboseq data (riboseq_l2fc and riboseq_padj; see Table S15).

**Table S13:** m^6^A modification of “EMT signature” genes The m^6^A-IP list was filtered against a list of known EMT signature genes. Ensemble gene ID (ENSG_ID); common gene name (gene_name); numerical m^6^A peak ID calculated by MACS2 (peak_ID); chromosome location (chr); chromosome coordinates for m^6^A peak start (start); chromosome coordinates for m^6^A peak end (end); location of the peak on the forward or reverse strand (strand); calculated log2foldchange and padj value of the gene (gene_l2fc and gene_padj); calculated log2foldchange of the m6A peak (peak_l2fc); peak_l2fc minus input gene_l2fc (diff_l2fc); padj values (DEseq_padj and logDEseqP); direction and significance of the padj and l2fc values (direction); presence and identity of the known RRACH motif for m^6^A modifications (m^6^A motif).

**Table S14:** Calculated half-life for UOK262-MUT and UOK262-WT genes

**Table S15:** Differential translation comparing UOK262-MUT and UOK262-WT and calculated using DEseq2

